# Bottlebrush polymers with sequence-controlled backbones for enhanced oligonucleotide delivery

**DOI:** 10.1101/2024.10.03.616318

**Authors:** Yun Wei, Peiru Chen, Mengqi Ren, Deng Li, Jiachen Lin, Tingyu Sun, Yuyan Wang, Shaobo Yang, Christopher Nenopoulos, Christopher Oetheimer, Yao Li, Chenyang Xue, Mona Minkara, Ke Zhang

## Abstract

The clinical translation of oligonucleotide-based therapeutics continues to encounter challenges in delivery. In this study, we introduce a novel class of delivery vehicles for oligonucleotides, which are based on polyethylene glycol (PEG) bottlebrush polymers with sequence-defined backbones. Using solid-phase synthesis and bespoke phosphoramidites, the oligonucleotide and the polymer backbone can both be assembled on the solid support. The synthesis allows chemical modifiers such as carbon 18 (C_18_) units to be incorporated into the backbone in specific patterns to modulate the cell-materials interactions. Subsequently, PEG side chains were grafted onto the polymer segment of the resulting polymer-oligonucleotide conjugate, yielding bottlebrush polymers. We report an optimal pattern of the C_18_ modifier that leads to improved cellular uptake, plasma pharmacokinetics, biodistribution, and antisense activity in vivo. Our results provide valuable insights into the pacDNA structure-property relationship and suggest a possibility of tuning the polymer backbone to meet the specific delivery requirements of various diseases.

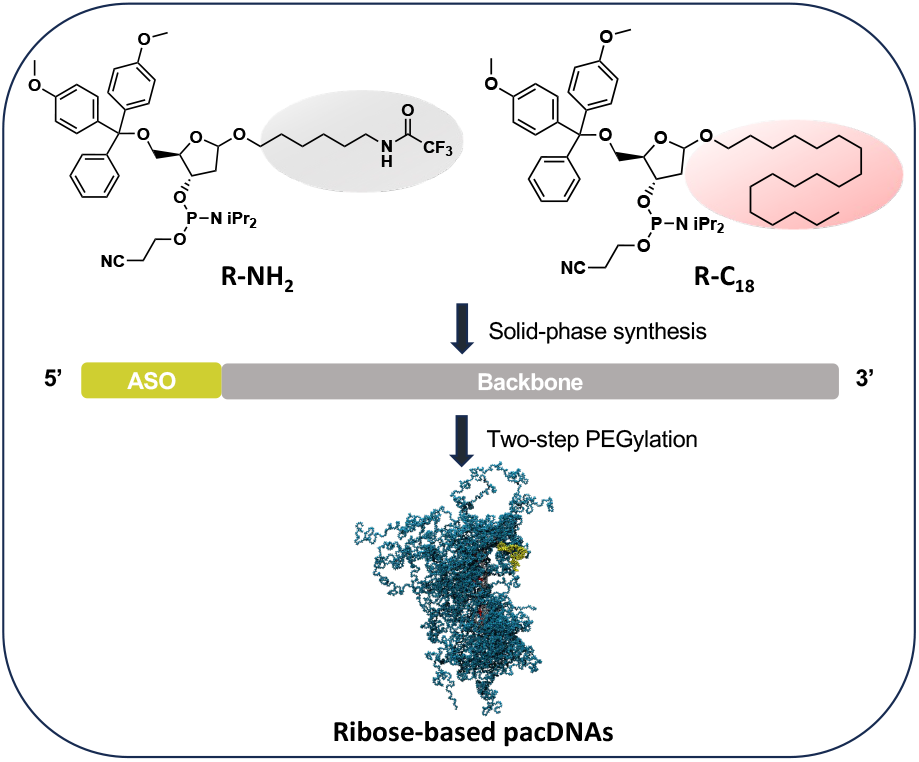

## Introduction

Oligonucleotide-based therapeutics hold immense promise for treating diseases through diverse mechanisms, such as gene regulation, receptor binding, and alternative splicing, among others.^1,2^ However, clinical development of oligonucleotide drug candidates often faces setbacks attributed to poor target engagement *in vivo* due to limited uptake by target organs and cells, limiting their use to a few concentrated disease settings.^3-5^ Current strategies to improve drug potency generally focus on chemical modifications, bioconjugation with antibodies, peptides, or small molecule ligands, and polyplex carriers such as lipid nano-particles and cationic polymers.^6-9^ However, concerns remain regarding potential toxicity and immunogenicity associated with chemical modifications and carriers, as well as suboptimal biodistribution following systemic administration.^10^ Thus, the development of a delivery system that can simultaneously enhance nuclease stability, facilitate rapid intracellular delivery of oligonucleotides, and ensure sufficient distribution into target tissues holds the potential to bridge the critical development gap.^9-11^

Previously, we have reported the design of a bottlebrush polymer-oligonucleotide conjugate (termed pacDNA), which effectively mitigates protein-oligonucleotide interactions while maintaining unaffected target RNA binding.^12,13^ This unique selectivity is achieved through the more densely packed poly(ethylene glycol) (PEG) side chains compared to typical linear or slightly branched PEG, which leads to reduced protein binding and consequently, fewer side effects *in vivo* that derive from unwanted protein-oligonucleotide interactions.^14^ Additionally, the large size of the conjugate reduces renal clearance and enhances circulation in the bloodstream, leading to significantly increased biodistribution to non-liver/kidney organs including difficult-to-target sites such as the skin, skeletal muscle, and heart.^15,16^

The bottlebrush polymer in our earlier system was prepared by ring-opening metathesis polymerization of norbornenyl monomers using Grubbs 3^rd^-generation catalyst. We identified three aspects where this system can be improved: 1) batch-to-batch consistency, 2) residue heavy metal content, and 3) control over the bottlebrush polymer backbone. Herein, we report the design, synthesis, and biological testing of a novel deoxyribose 3’-5’ phos-phodiester-derived bottlebrush-oligonucleotide conjugate, which offers unprecedented flexibility and control in the backbone chemistry, length, sequence, and composition. We demonstrate an optimized backbone structure containing patterned C_18_ units for application in targeting Kirsten Rat Sarcoma Virus (KRAS) mRNA in a GEMM-derived mouse allograft model. These findings provide valuable insights for the development of tailored vehicles to meet indication-specific delivery requirements.

## Results and Discussions

The poly(2-deoxyribose phosphodiester) backbone polymer is constructed using a stepwise condensation approach, employing two bespoke modified phosphoramidites: ribose-NH_2_ (R-NH_2_) and ribose-C_18_ (R-C_18_). The modifiers were synthesized from 1-chloro-3,5-di(4-chlor-benzoyl)-2-deoxy-D-ribose in ∼40% overall yields in multigram scales (**Figures 1A, S10-25; Schemes S1, S2**). The oligonucleotide component was synthesized as an integral part of the backbone on the solid support.

**Figure 1.**
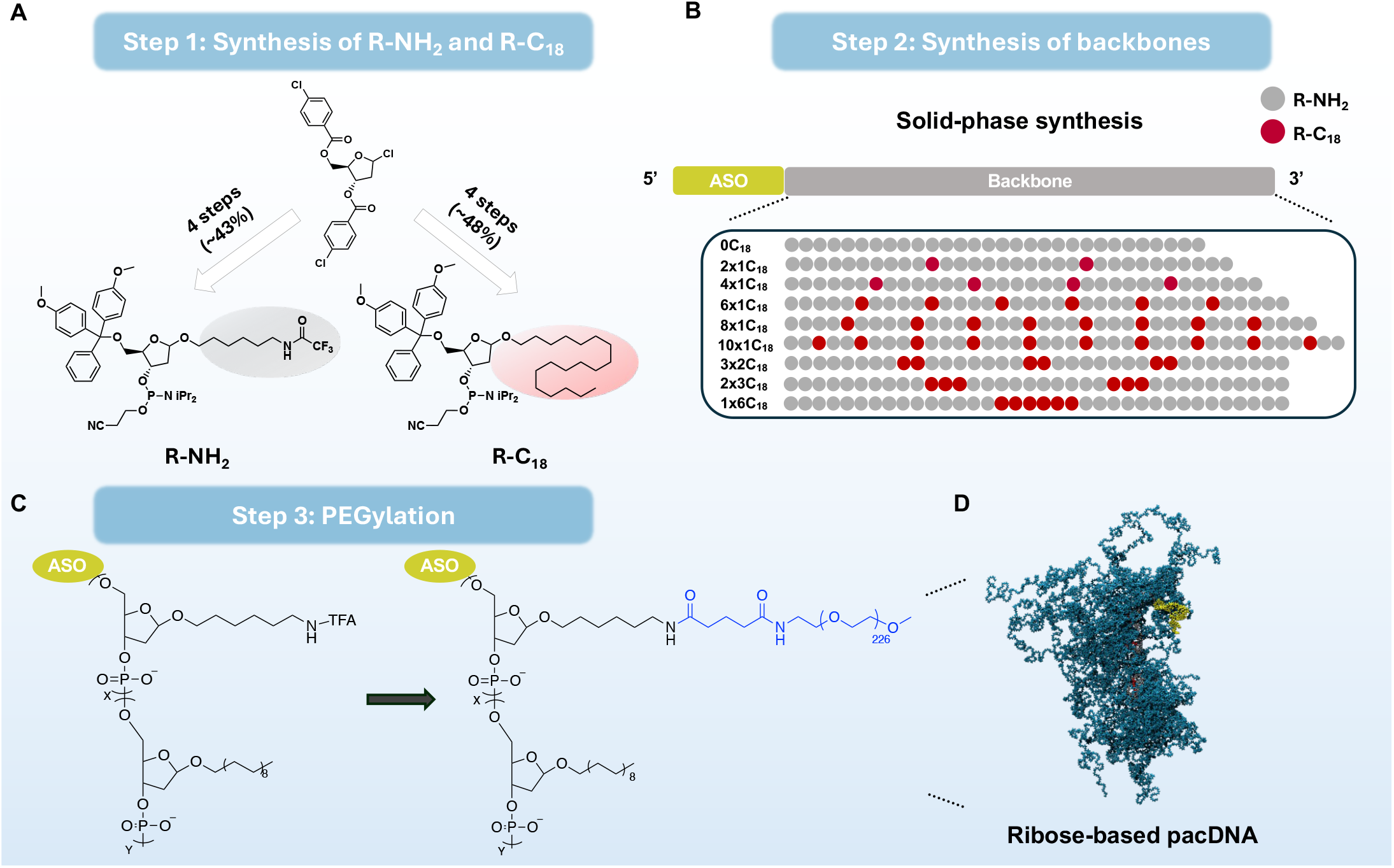
Design of ribose-based pacDNAs. (**A-B**) Synthetic scheme and backbone designs of pacDNAs. The grey and red circle represents R-NH_2_ unit and R-C_18_ unit, respectively. ASO: antisense oligonucleotide. (**C**) Schematics of PEGylation process of ribose-based pacDNA. (**D**) Molecular dynamics simulation of ribose-based pacDNA with four C_18_ modifiers (cyan: PEG; grey: backbone; red: C_18_; yellow: ASO).

Due to the utilization of the same phosphoramidite chemistry, the oligonucleotide can be positioned at either the 3’ or the 5’ of the polymer backbone without requiring additional post-conjugation steps. For the purposes of this proof-of-concept study, antisense oligonucleotides (ASOs) targeting the 3’ untranslated region (UTR) of both human and mouse KRAS mRNA (ASO1 and ASO2, respectively) were selected as the payload for this ribose-based pacDNA, which were positioned at the 5’ of the ribose backbone **(Figure 1B)**.^17,18^ The backbone is designed to incorporate 30 repeating units of ribose-NH_2_, which can be subsequently PEGylated to give the bottlebrush morphology **(Figure 1C, 1D)**.^16,19^ Because the phosphodiester ribose backbone is significantly more hydrophilic than the polynorbornene (PN)-based pacDNA, which may limit materials-cell membrane interactions leading to reduced cell uptake, we investigated how hydrophobic C_18_ modifiers introduced into the backbone can affect the cellular uptake and biodistribution *in vivo*. We varied the numbers of C_18_ modifiers (ranging from 0 to 10) as well as their distribution patterns (for backbones containing six C_18_ modifiers) in order to probe the structure-property relationship **(Figure 1B)**.

Following the completion of solid-phase backbone/ASO synthesis, the trifluoroacetyl protecting groups were removed and the hybrid strand was cleaved from the solid support. The successful strand was isolated using a dimethoxytrityl (DMT)-affinity column. To construct the bottlebrush structure, the amine groups on the backbone were derivatized with a heterodifunctional N-hydroxy-succinimide (NHS)- and methyl-terminated 10kDa PEG, using a two-stage process.^19^ The purified strand was PEGylated initially in 1× phosphate buffered saline (PBS) at 4 °C overnight. The partially PEGylated product was desalted, lyophilized, and subsequently reacted with another equivalent of PEG in anhydrous *N, N*-dimethylformamide (DMF) for full derivatization. Excess PEG and residues were removed by aqueous gel permeation chromatography (GPC), yielding highly uniform, narrowly dispersed pacDNA structures, as evidenced by aqueous and DMF GPC **(Figure 2A, Table S2)**, dynamic light scattering (DLS) **(Figure 2B, S1)**, and transmission electron microscopy (TEM) (**Figure 2C**). The uniformity of pacDNA particles were consistently observed across all samples irrespective of the quantity or arrangement of R-C_18_ units, suggesting that the C_18_ modifiers do not cause aggregation in solution. ζ potential measurement revealed that all pacDNA samples are slightly anionic (−3.26mV to -11.71mV) in Nano-pure™ water compared with free oligonucleotide (− 33.6mV) **(Figure 2D)**. Collectively, these results demonstrate that ribose-based pacDNAs can be robustly synthesized, allowing for fine tuning of the size/backbone sequence of the pacDNAs and the potential to modulate their bioactivities. A comparison between ribose- and PN-based pacDNAs can be found in **Figure S2**.

**Figure 2.**
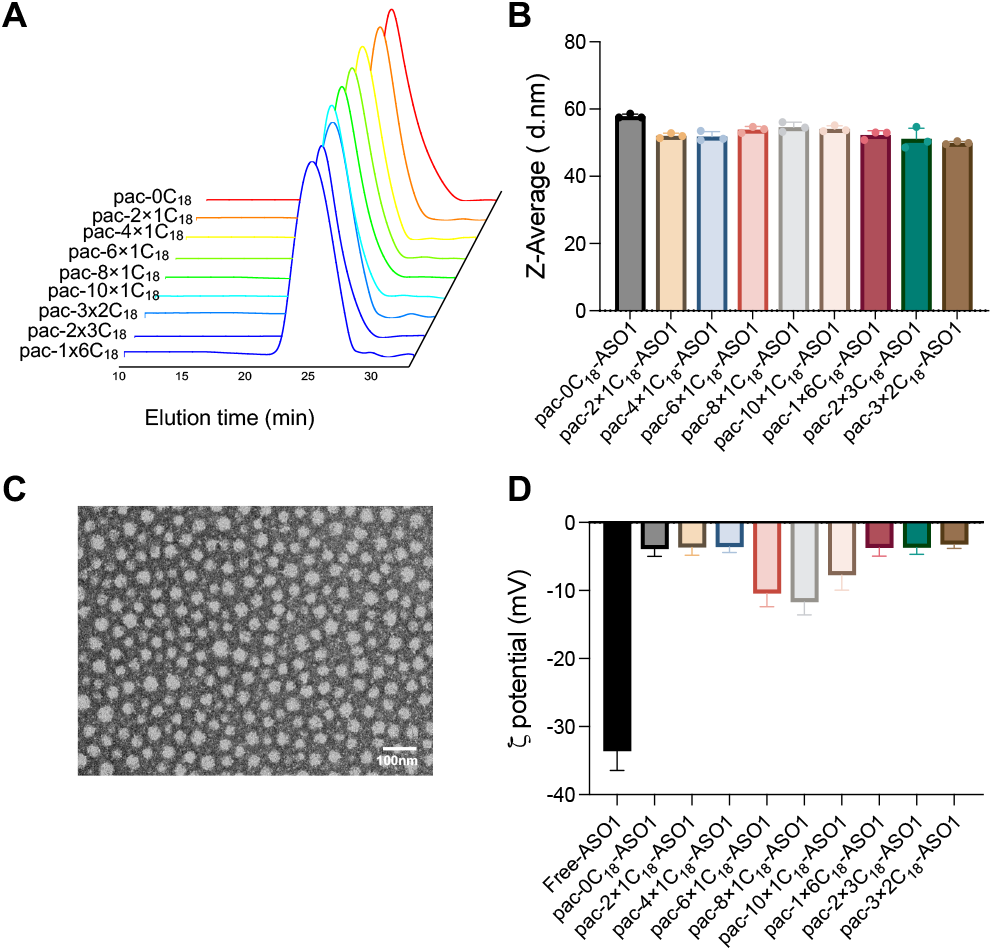
Characterization of ribose-based pacDNAs. (**A**) Aqueous GPC chromatograms of pacDNAs after two-stage PEGylation. (**B**) pacDNAs Z-average molecular size in Nano-pure™ water as determined by DLS. (**C**) Representative TEM images of pacDNAs (pac-4×1C_18_-ASO1) with negative staining using 2% uranyl acetate. (**D**) pacDNAs ζ potential in Nano-pure™ water.

Next, we assessed the intracellular delivery efficacy of the pacDNA panel in NCI-H358 cells, a non-small lung carcinoma line harboring KRAS^G12C^ mutation. Flow cytometry analysis revealed that as the number of R-C_18_ units increased within the bottlebrush backbone, cellular uptake also increased **(Figure 3A)**, suggesting the structurally more hydrophobic pacDNA constructs may exhibit stronger materials-cell interactions. The trend does not change when bottlebrush polymers lacking the ASO component were tested (**Figure S3A**). Interestingly, when the same number of C_18_ units were incorporated into the bottlebrush backbone, the evenly distributed patterns demonstrated higher cellular uptake efficiencies **(Figure 3B)**, while the clustering of C_18_ units reduced cell uptake. In fact, when all six C_18_ units were clustered together (pac-1×6C_18_-ASO1), the cellular uptake was comparable to that of the pacDNA without any C_18_ units (pac-0C_18_-ASO1). The flow cytometry measurements were further supported by confocal microscopy. Cy5-labeled pacDNAs were incubated with NCI-H358 cells for 8 h before imaging. Again, pacDNAs with a higher number of C_18_ units showed greater cell uptake (**Figure 3C, S4**). We postulate that when R-C_18_ units are positioned adjacent to one another, the self-interaction among the C_18_ decreases their tendency to interact with the cellular membrane. In contrast, separating the C_18_ units spatially reduces such self-interactions, leading to a higher potential energy state and stronger tendency to bind with cell membrane upon contact. Increased cellular uptake of ASO by lipid conjugation has been reported.^20,21^ However, lipid conjugates often exhibit increased cytotoxicity, possibly due to membrane-lytic activity of the amphiphilic conjugate.^22,23^ In contrast, the pacDNAs do not cause cytotoxicity (**Figure S3B**) even though they contain multiple C_18_ units. One interpretation is that the sterics of the pacDNA make it difficult for the lipid chains to aggregate and exhibit surfactant-like properties, limiting potential membrane lytic activity.

**Figure 3.**
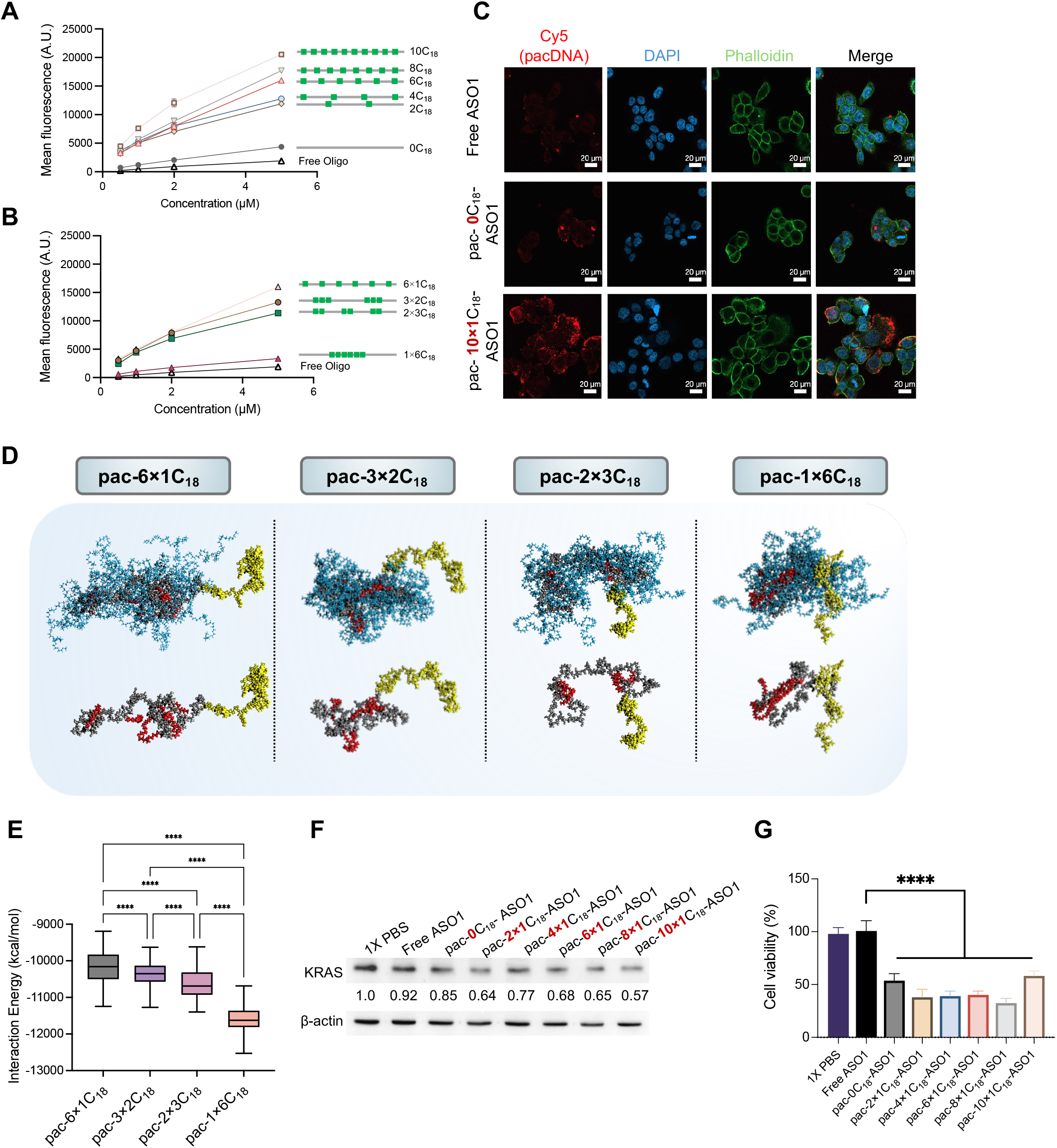
*In vitro* properties of ribose-based pacDNAs. (**A-B**) Cellular uptake by NCI-H358 cells of Cy5-labeled pacDNAs-ASO1 containing varying numbers and arrangement of C18 modifiers after 4 h incubation, as determined by flow cytometry. MFI: mean fluorescence intensity. (**C**) Confocal microscopy of NCI-H358 cells treated with Cy5-labeled free ASO1 or pacDNAs-ASO1 for 8h (additional samples: Figure S4). (**D-E**) Interaction energy of pacDNAs after interaction with water as determined by molecular dynamics simulation and representative molecular snapshots from the simulation trajectory (up row is the structures of entire molecules, down row is the corresponding structures without PEG chains to show backbones more clearly; cyan: PEG; grey: backbone; red: C18; yellow: ASO). (**F**) KRAS depletion efficiency of 10μM pacDNAs-ASO1 in NCI-H358 cells. The number shows relative KRAS protein level analyzed by ImageJ software. (**G**) Inhibitive effect of 10μM pacDNAs-ASO1 on the proliferation of NCI-H358 cells after 48 h incubation.

To gain molecular level insights, we carried out full atomistic molecular dynamics (MD) simulations to quantitatively assess the interaction energy between water molecules and pacDNAs with six C_18_ units arranged in different distribution patterns. In this context, interaction energy refers to the total energy arising from all intermolecular forces between the pacDNA molecules and surrounding water molecules. A more negative interaction energy indicates stronger interactions with water, leading to greater solubility of the pacDNA in aqueous environments. We hypothesize that the pacDNA with the least amount of lipid self-interactions will be the most prone to membrane binding. This hypothesis can be tested by measuring pacDNA interaction energy in water – structures with the least interaction energy with water (most difficult to dissolve) should exhibit the highest cell uptake. To make the simulation computationally feasible, all PEG side chains were truncated to a 20-mer, which was found to be sufficiently large to inhibit backbone coiling under simulation conditions. The simulation trajectory confirms more extensive lipid-lipid interactions for clustered vs. spaced arrangement of C_18_ units (**Figure 3D**). The pacDNA with the smallest spacing (pac-1×6C_18_) exhibited the most interaction energy (most negative). This result suggests that clustering of C_18_ units increases solubility in water and therefore reduces C_18_ interactions with cell membranes (**Figure S3E**). Conversely, the pacDNA with the largest spacing (pac-6×1C_18_) exhibited the lowest interaction energy (least negative), indicating that pac-6×1C_18_ has the least tendency to be dissolved in water, and by extension, an enhanced propensity for interacting with cell membrane and cellular uptake. These observations underscore the pivotal role of the digital backbone design, which allows for precise tuning of the biophysical behavior of pacDNAs.

Next, we assessed the efficacy of the pacDNAs to engage with cytosolic mRNA targets using pacDNAs bearing ASO1 (*vide supra*) and varying numbers of evenly distributed C_18_ units. Western blot analysis showed the downregulation of KRAS protein levels in all pacDNA-treated groups (**Figure 3F**). Interestingly, when the pacDNA has at least two C_18_ units, KRAS depletion levels did not vary significantly. We also analyzed cell viability using a 3-(4,5-dimethylthiazol-2-yl)-2,5 diphenyl tetrazolium bromide (MTT) assay. All pacDNAs significantly inhibited tumor cell growth, with inhibition levels ranging from 40% to 60% compared to non-treated cells while the free ASO1 showed no inhibitory activity (**Figure 3G, S5**).

Prior to nominating a candidate for efficacy studies, it is important to understand how the pacDNA backbone series affects plasma pharmacokinetics (PK) and biodistribution. Cy5-labeled pacDNAs were intravenously (i.v.) administered to C57BL/6 mice via the tail vein, and blood plasma samples were collected by submandibular puncture at pre-determined time points over a 72-hour period. Compared to free ASO, all pacDNA constructs, regardless of the number of R-C_18_ units in the backbone, exhibited significantly prolonged circulation times (**Figure 4A**). Analyzing the plasma PK using a two-compartment model, it can be seen that the distribution half-lives (t_1/2α_) generally decrease with increasing R-C_18_ number (**Figure 4B**). Interestingly, a straightforward correlation was not observed for the elimination half-life (t_1/2β_), which increases first and then decreases with C_18_ numbers, peaking at pac-4×1C_18_. The insertion of two or four R-C_18_ units may represent optimized structures because these specific arrangements are capable of significantly enhancing the bioavailability of the ASO as characterized by area-under-the-curve (AUC_∞_), while being sufficiently strong for cellular uptake.

**Figure 4.**
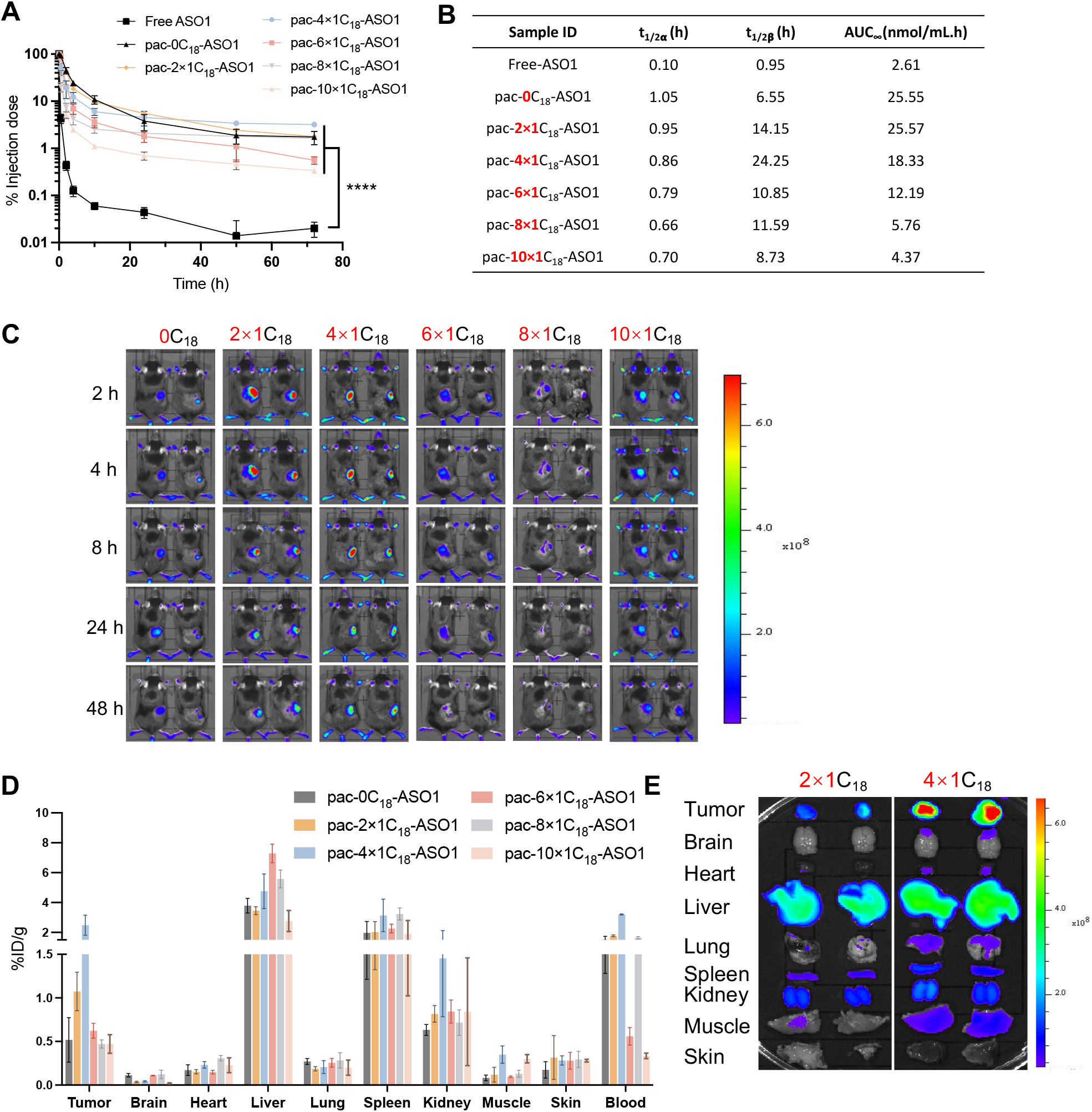
Plasma pharmacokinetics and biodistribution of ribose-based pacDNAs. (**A**) Plasma pharmacokinetics of Cy5-labeled pacDNA and free DNA in C57BL/6 mice following i.v. injection. (**B**) Plasma pharmacokinetics parameters of ribose-based pac-DNAs with variant numbers of C_18_ modifier. (**C**) Live animal fluorescence imaging of C57BL/6 mice bearing K273 allograft following i.v. injection of Cy5-labeled pacDNA. Areas surrounding the allograft has been shaved to facilitate imaging. (**D-E**) Quantitative biodistribution and *ex vivo* imaging of pacDNAs in major organs/tissues of tumor-bearing C57BL/6 mice 72 h post i.v. injection.

We next carried out a biodistribution study comparing the different pacDNA constructs using C57BL/6 mice bearing K273 tumor allograft. The K273 cell is a KRAS^G12D^ lung cancer line derived from a genetically engineered mouse model (GEMM). The pacDNAs were administered i.v., and fluorescence images were captured at predetermined time points (**Figure 4C**). The fluorescence signal at the tumor site first increases and then decreases with increasing C_18_ numbers, following a similar trend of the plasma elimination half-life. To quantitatively analyze pacDNA biodistribution among major organs and tissues, mice were sacrificed 72 hours after injection, and tissue lysates were used to determine distribution as % injected dose per gram of tissue (%ID/g, **Figure 4D**). Again, pacDNAs with two or four R-C_18_ units exhibit stronger tumor localization compared to those with six, eight, or ten C_18_ modifiers. Because pac-4×1C_18_ shows the highest tumor distribution compared with other backbone designs (**Figure 4E, S6**), we focused on this specific arrangement for subsequent studies.

To investigate the antitumor efficacy of pac-4×1C_18_ using the K273 allograft model, we prepared pac-4×1C_18_-ASO2, which is matched to the mouse wildtype KRAS mRNA sequence at the 3’ UTR region. First, we validated the potency of pac-4×1C_18_-ASO2 using the K273 cell line *in vitro*. Western blot shows that pac-4×1C_18_-ASO2 downregulated KRAS protein levels in a dose-dependent manner (**Figure S7A**). MTT cell viability assay showed that pac-4×1C_18_-ASO2 reduced K273 cell viability, while free ASO2 or pac-4×1C_18_-scramble showed no inhibitory effect (**Figure S7B)**. The antitumor efficacy of pac-4×1C_18_-ASO2 was assessed in female C57BL/6 mice bearing subcutaneous K273 allografts. When the allografts reached a volume of 100 mm^3^, 0.5 µmol/kg of pac-4×1C_18_-ASO2, free ASO2, or vehicle (PBS) were administrated i.v. once every four days for a total of four doses (**Figure 5A**). By day 24 after tumor inoculation, the average tumor volume of vehicle group reached 864 mm^3^, while pac-4×1C_18_-ASO2 significantly inhibited tumor growth and extended mice survival with the average tumor volume at 299 mm^3^ (**Figure 5B, 5C**). In contrast, free ASO2 elicited no survival benefit. Neither pac-4×1C_18_-ASO2 nor free ASO2 lead to significant body weight changes (**Figure S8**). Immunohistostaining of tumor tissue revealed reduced KRAS protein expression after pac-4×1C_18_-ASO2 treatment but not with free ASO2 (**Figure 5D**).

**Figure 5.**
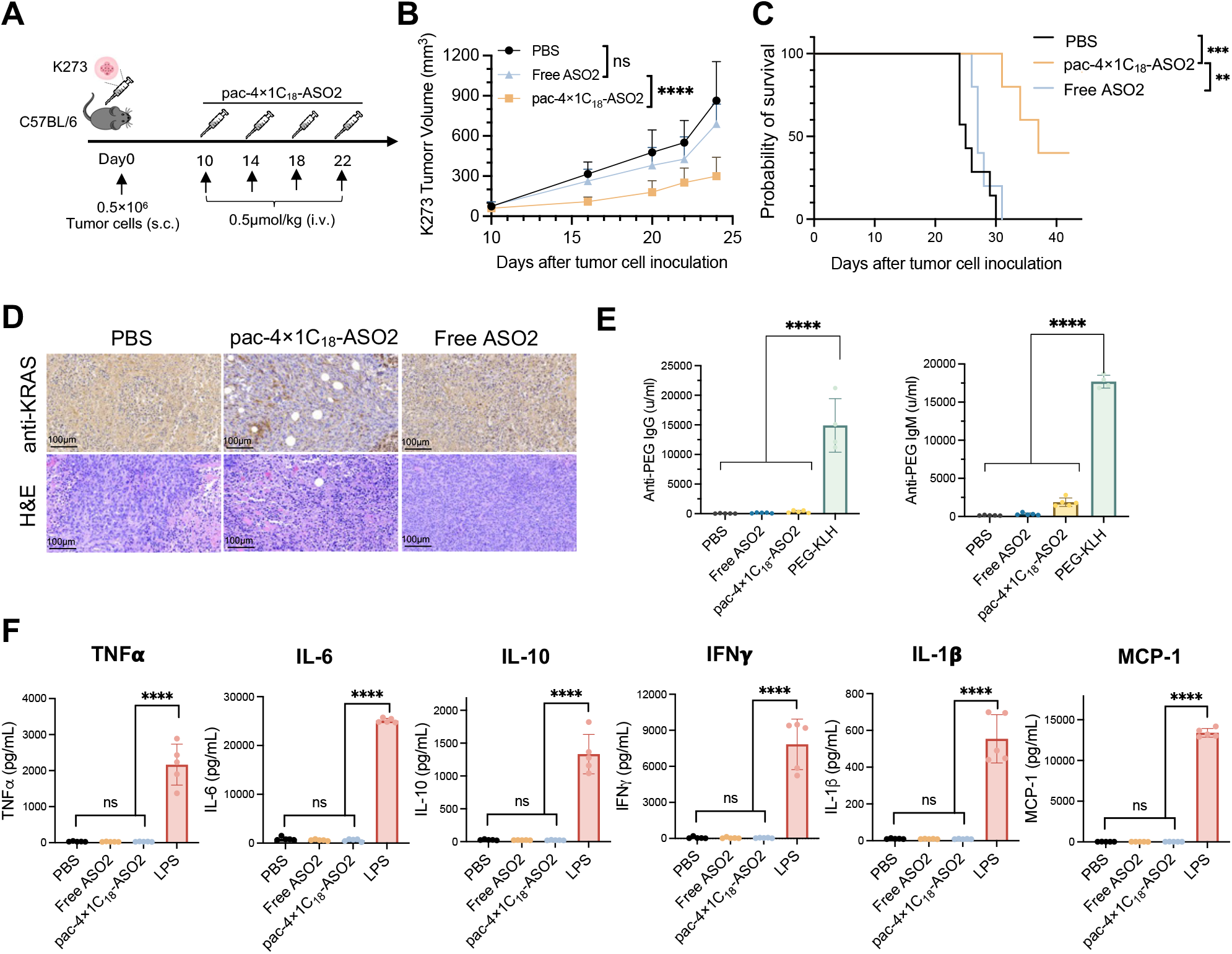
*In vivo* antitumor efficacy of ribose-based pacDNAs. (**A**) K273 allograft volume changes in 24 days with i.v. injection of PBS, ASO2, and pac-4×1C_18_-ASO2 at equivalent ASO doses (0.5 µmol/kg) every four days since day 10. Injections are indicated by black arrows. Statistical significance was performed using two-way ANOVA. (**B**) Mean tumor growth curve. (**C**) Kaplan-Meier endpoint animal survival analysis. Statistical analysis was calculated by the log-rank test. (**D**) Immunohistostaining and H&E staining of K273 tumor after treatment. (**E**) Anti-PEG IgM and IgG levels in C57BL/6 mice plasma after i.v. injection of PBS, free ASO2, pac-4×1C_18_-ASO2 and PEG-KLH (positive control) at 0.5 µmol/kg every four days for four doses. Plasma was collected one week after the last dose. Statistical analysis was performed using one-way ANOVA. (**F**) Selected cytokine and chemokine levels in C57BL/6 mice plasma 5 h after i.v. injection of PBS, free ASO2, pac-4×1C_18_-ASO2 (2 µmol/kg), and LPS (0.5 mg/kg). Statistical analysis was performed using one-way ANOVA. ^****^ *P* <0.0001, ^***^ *P* <0.001, ^**^ *P* <0.01, ^*^ *P*< 0.05.

Next, we performed an *in vivo* safety analysis of pac-4×1C_18_-ASO2. Hematoxylin and eosin (H&E) staining demonstrated no obvious histological changes in major internal organs (**Figure S9**). Macromolecular therapeutics requiring repeated dosing may induce anti-drug adaptive immunity, leading to loss of drug activity in subsequent doses. To evaluate the anti-PEG immunoglobulin IgM/IgG response, healthy C57BL/6 mice were administrated pac-4×1C_18_-ASO2 or free ASO2 every four days for four doses. Mouse plasma was collected one week after the last dose. Enzyme-linked immunosorbent assays (ELISA) revealed that pac-4×1C_18_-ASO2 induced very limited levels of anti-PEG IgM/IgG (**Figure 5E**). Both responses were minor compared to positive control (PEG-keyhole limpet hemocyanin [KLH] conjugate). To further investigate possible unintended activation of the immune system, C57BL/6 mouse plasma was collected 3 h after i.v. injection of pac-4×1C_18_-ASO2 or free ASO2, and cytokines and chemokines related to innate and adaptive immune responses were measured. The assay detected no apparent cytokines or chemokines for either pac-4×1C_18_-ASO2 or free ASO2 treatment groups, while lipopolysaccharide (LPS) i.v. injection (positive control) generated high levels of cytokines/chemokines (**Figure 5F**).

## Conclusion

In conclusion, our study presents a robust approach to construct bottlebrush polymers with arbitrary control of the backbone size, composition, and monomer sequence. These capabilities allow us to take the advantage of C_18_ modifiers and optimize the biological properties such as cell uptake and tumor localization. Experimental and computational studies suggest that the different arrangements of C_18_ affect their self-interaction, which in turn modifies their readiness to engage with the cell membrane. Through optimization, we identified pac-4×1C_18_ as overall the most favorable given *in vitro* cell uptake, gene silencing, and biodistribution in mice. In a KRAS^G12D^ lung cancer allograft GEMM mouse model, a KRAS-depleting pac-4×1C_18_ showed single-agent tumor suppressive potency at a fraction of concentration typically used for the ASO modality. Taken together, our results provide valuable insights into the pacDNA structure-property relationship and suggest a possibility of tuning the polymer backbone to meet the delivery requirements of various diseases.

## ASSOCIATED CONTENT

### Supporting Information

Materials, experimental procedures and methods, sequences, synthesis, NMR spectroscopy, DLS, dynamic simulation, confocal, in vitro assays, and H&E staining of mice major organs after treatment. This material is available free of charge via the Internet at http://pubs.acs.org.

## Author Contributions

Y.W., P.C., and M.R. contributed equally. K.Z., P.C., Y.W. and M.R. conceived the project, designed the experiments, and analyze the data. Y.W. and P.C. completed sample synthesis the biological studies. M.R. designed synthesis routes and analyzed NMR results. D.L. and M.M. contributed to the dynamic simulation studies. S.Y. contributed to confocal studies. All other authors contributed to material synthesis. K.Z., P.C. and Y.W. cowrote the manuscript. The manuscript was written through contributions of all authors. All authors have given approval to the final version of the manuscript.

## Funding Sources

This publication was made possible by the National Science Foundation (DMR award number 2004947), the National Institute of General Medical Sciences (1R01GM121612), and the National Cancer Institute (4R42CA275425 and 5R01CA251730).

### Notes

K.Z. hold financial interest in pacDNA Inc., a company commercializing the pacDNA technology.

## ACKNOWLEDGMENT

We gratefully acknowledge Dr. Jean Zhao for gifting K273 cell line. We are grateful to the Northeastern University-Institutional Animal Care and Use Committee for supporting our animal studies. We thank Dr. Guoxin Rong from the Institute for Chemical Imaging of Living System at Northeastern University for assistance with confocal microscopy. We appreciate Rafay Abu from Barnett Institute of Chemical and Biological Science for HRMS analysis.

