## Supporting Information for "Bottlebrush polymers with sequence-controlled backbones for enhanced oligonucleotide delivery"

### for

#### Materials and reagents

1-Chloro-3,5-di(4-chlorbenzoyl)-2-deoxy-D-ribose, 6-(Trifluoroacetamido) hexanol, 4,4′-Dimethoxytrityl chloride, 1-Octadecanol, and 2-Cyanoethyl N,N-diisopropylchlorophosphoramidite were purchased from Sigma-Aldrich Co., USA. Methoxy polyethylene glycol (PEG) glutaramide succinimidyl ester ($M_n$=10 kDa) was purchased from Creative PEGWorks (Chapel hill, NC, USA). Phosphoramidites and supplies for DNA synthesis were obtained from Glen Research Co., USA. All other materials were obtained from Fisher Scientific Inc., USA, or VWR International LLC., USA, and were used as received unless otherwise indicated.

#### Instrumentations

$^1$H and $^{13}$C nuclear magnetic resonance (NMR) spectra were recorded on a Bruker 700 MHz NMR spectrometer (Bruker Scientific LLC, MA, USA). Reversed-phase high-performance liquid chromatography (RP-HPLC) was performed on a Waters (Waters Co., MA, USA) Breeze 2 HPLC system coupled to a Symmetry® C18 3.5 µm, 4.6×75 mm reversed-phase column and a 2998 PDA detector, using TEAA buffer (0.1 M) and HPLC-grade acetonitrile as mobile phases. Aqueous GPC measurements were performed on a Waters Breeze 2 GPC system equipped with Ultrahydrogel™ 1000, Ultrahydrogel™ 500 and Ultrahydrogel™ 250, 7.8Å×300 mm column and a 2998 PDA detector (Waters Co., MA, USA). Phosphate-buffered saline (PBS, pH 7.4) was used as the eluent running at a flow rate of 0.8 mL/min. *N, N*-dimethylformamide (DMF) GPC analysis was performed on EcoSEC HLC-8320 GPC system (Tosoh Bioscience LLC, Tokyo, Japan) equipped with a TSKGel α-M 7.8×300 mm, 13 µm column and RI/UV-Vis detectors. The mobile phase is 0.05 M lithium bromide in HPLC-grade DMF, and samples were analyzed at a flow rate of 0.4 mL/min. DMF GPC calibration was based on a ReadyCal kit of polyethylene glycol standards (PSS-Polymer Standard Service-USA Inc., MA, USA). The kit covers an $M_n$ range from 232 Da to 1015 kDa. For transmission electron microscopy (TEM) measurements, samples were imaged on a JEOL JEM 1010 electron microscope with an accelerating voltage of 80 kV. High-resolution mass spectra (HRMS) analysis was performed on a Q Exactive Orbitrap (Thermofisher Scientific™ ) using a syringe pump with direct infusion. Data was acquired in positive/negative ion mode using a hybrid orbitrap Q Exactive mass spectrometer (Thermo Fisher Scientific, Bremen, Germany) with direct infusion electrospray ion source (HESI) using a syringe pump. The flow rate was 5ul/min with +4.5 kV /-3.5 kV electrospray voltages. Analysis was performed in full scan mode with a mass range of 150-2000 m/z, including an automatic gain control setting of $1\times10^6$ and a mass resolution of 17,000. DLS and ζ potential measurements were performed on a Malvern Zetasizer Nano-ZSP (Malvern, UK). Samples were dissolved in Nanopure™ water at a concentration of 1 µM and filtered through a 0.2 µm PTFE filter before measurement. Histopathology (H&E) and immunohistochemistry studies were carried out at iHisto Inc.

**Scheme S1. Synthesis of ribose-amine phosphoramidite (R-$NH_2$)**

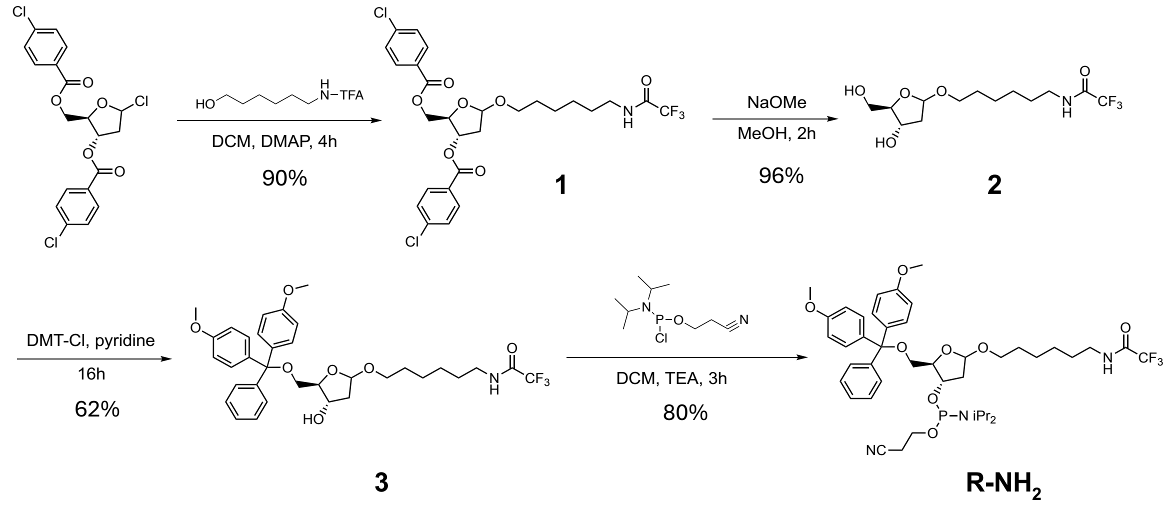


**Synthesis of TFA-amine-ribose (1, Scheme S1)** 1-Chloro-3,5-di(4-chlorbenzoyl)-2-deoxy-D-ribose (1.04g, 2.42 mmol, 1 eq.) was dissolved with 10 mL of anhydrous DCM in a round bottom flask. 6-(Trifluoroacetamido) hexanol (0.62 g, 2.90 mmol, 1.2 eq.) and 4-Dimethylaminopyridine (DMAP) (87.90 mg, 0.72 mmol, 0.3 eq.) were dissolved in 5 mL of anhydrous DCM, separately, and the solutions were added into the flask. The reaction was allowed to react at room temperature (R.T.) for at least 4 h. Afterwards, the reaction was concentrated by rotary-evaporation and purified by flash chromatography on silica column using a dichloromethane with ethyl acetate (0 to 5%) eluent system. The product was then dried under high vacuum to yield a white solid. **$^1$H NMR (700 MHz, $CDCl_3$)** δ 7.93 – 7.86 (m, 4H), 7.37 – 7.31 (m, 4H), 6.24 (s, 1H), 5.33 (ddd, *J* = 8.0, 3.7, 2.2 Hz, 1H), 5.22 – 5.19 (m, 1H), 4.53 (dd, *J* = 11.7, 4.0 Hz, 1H), 4.48 – 4.40 (m, 2H), 3.66 (dt, *J* = 9.6, 6.7 Hz, 1H), 3.37 (dt, *J* = 9.6, 6.3 Hz, 1H), 3.25 (q, *J* = 6.9 Hz, 2H), 2.42 (ddd, *J* = 14.6, 8.0, 5.3 Hz, 1H), 2.13 (ddd, *J* = 14.4, 2.2, 1.0 Hz, 1H), 1.56 – 1.42 (m, 5H), 1.38 – 1.23 (m, 4H). **$^{13}$C NMR (176 MHz, $CDCl_3$)** δ 165.46, 165.40, 157.27, 157.06, 139.81, 139.70, 131.16, 131.13, 131.04, 128.82, 128.77, 128.21, 116.68, 115.05, 103.79, 80.75, 75.09, 67.40, 64.59, 39.88, 39.11, 29.52, 28.96, 28.85, 26.42, 25.75. **HRMS**: calculated for $C_{27}H_{28}Cl_2F_3NO_7$ [M-H]$^-$: 604.12, found: 604.11.

**Synthesis of deprotected TFA-amine-ribose (2, Scheme S1)** Compound **1** (1.01 g, 1.66 mmol, 1 eq.) was dissolved in 9 mL 0.5 M NaOMe in methanol (4.16 mmol, 2.5 eq.) and allowed to stir at R.T. for 2 h. The solution was dried loaded on silica column and purified using a 5% ethyl acetate in dichloromethane added with methanol (0 to 5%) eluent system. The deprotected compound **1** was then dried under high vacuum to yield a viscous transparent liquid (Compound **2**). **$^1$H NMR (700 MHz, $CDCl_3$)** δ 7.42 (t, *J* = 5.9 Hz, 1H), 5.19 – 5.13 (m, 1H), 4.43 (td, *J* = 6.2, 3.6 Hz, 1H), 4.13 – 4.03 (m, 1H), 3.99 (q, *J* = 3.9 Hz, 1H), 3.73 – 3.65 (m, 2H), 3.65 – 3.55 (m, 2H), 3.38 (tt, *J* = 9.7, 5.7 Hz, 2H), 3.34 – 3.25 (m, 2H), 2.21 (ddd, *J* = 13.8, 7.0, 2.1 Hz, 1H), 2.15 – 2.04 (m, 1H), 1.54 (dtd, *J* = 15.7, 12.4, 7.2 Hz, 4H), 1.32 (tq, *J* = 14.6, 7.3 Hz, 4H). **$^{13}$C**

**NMR (176 MHz, $CDCl_3$)** δ 157.81, 157.60, 157.58, 157.39, 157.37, 157.18, 116.75, 115.11, 104.41, 87.22, 71.93, 68.00, 63.53, 42.23, 39.77, 29.27, 28.61, 26.24, 26.20, 25.56. **HRMS**: calculated for $C_{13}H_{22}F_3NO_5$ [M-H]$^-$: 328.15, found: 328.14.

**Synthesis of TFA-amine-ribose-DMT (3, Scheme S1)** Compound **2** (0.55 g, 1.65 mmol, 1 eq.) was dissolved in 10 mL anhydrous pyridine and the solution was allowed to cool with an ice-bath. Dimethoxytrityl chloride (0.62 g, 1.82 mmol, 1.1 eq.) was dissolved in 5 mL anhydrous pyridine and added dropwise into the reaction under positive nitrogen flow. The reaction was allowed to gradually reach to R.T. and stirred overnight. Afterwards, the solution was quenched with methanol and concentrated by rotary evaporation to remove pyridine using toluene to co-evaporate. The concentrated reaction was then dissolved in ethyl acetate and extracted with 0.1 M $NaHCO_3$ (2x) and brine (2x). The final solution was dried with $MgSO_4$, concentrated by rotary evaporation, and purified by flashing chromatography on silica column with a hexane: ethyl acetate (3:1 to 1:1, v:v) eluent system supported with 0.1% triethylamine. The product was then dried under high vacuum to yield viscous liquid, orange in color. **$^1$H NMR (700 MHz, $CDCl_3$)** δ 7.34 – 7.10 (m, 9H), 6.73 (d, *J* = 8.7 Hz, 4H), 5.14 (d, *J* = 4.5 Hz, 1H), 4.13 (dt, *J* = 5.4, 2.8 Hz, 1H), 4.09 (dd, *J* = 9.4, 5.8 Hz, 1H), 3.67 (s, 7H), 3.35 – 3.29 (m, 1H), 3.27 – 3.18 (m, *J* = 6.7 Hz, 2H), 3.08 – 2.95 (m, 3H), 2.10 (ddd, *J* = 13.6, 6.0, 4.6 Hz, 1H), 1.95 – 1.90 (m, 1H), 1.54 – 1.44 (m, 4H), 1.27 (qd, *J* = 9.5, 5.7 Hz, 4H). **$^{13}$C NMR (176 MHz, $CDCl_3$) δ** 158.49, 157.39, 157.19, 144.83, 136.03, 135.93, 130.07, 128.16, 127.84, 126.80, 116.77, 115.14, 113.14, 104.35, 86.80, 86.12, 73.54, 67.22, 64.05, 55.21, 40.89, 39.82, 29.42, 28.83, 26.33, 25.77. **HRMS**: calculated for $C_{34}H_{40}F_3NO_7$ [M-H]$^-$: 630.28, found: 630.27.

**Synthesis of ribose-amine (R-$NH_2$, Scheme S1)** Compound **3** (0.42 g, 0.65 mmol, 1 eq.) was dissolved in 5 mL anhydrous dichloromethane and 452.9 µL of triethylamine (0.33 g, 3.25 mmol, 5 eq.) was added to the solution. The solution was cooled with an ice-bath and undergo continuous nitrogen flow. The 2-Cyanoethyl N, N-diisopropylchlorophosphoramidite was dissolved in 2 mL anhydrous dichloromethane and added into the previous reaction mixture. The reaction was allowed to stir on ice for 30 min and at R.T. for 2.5 h. Ethyl acetate was added to dilute the reaction and extracted with 0.1 M $NaHCO_3$ (2x) and brine (2x). The solution was dried with $MgSO_4$ and concentrated via rotary-evaporation. The mixture was purified by flashing chromatography on silica column with a hexane: ethyl acetate (2:1 to 1:1, v:v) eluent system supported with 0.1% triethylamine. The final product was dried under high vacuum yielding a viscous liquid. **$^1$H NMR (700 MHz, $CDCl_3$)** δ 7.39 – 7.18 (m, 9H), 6.78 – 6.72 (m, 4H), 5.14 (dd, *J* = 5.4, 1.6 Hz, 1H), 4.28 (ddt, *J* = 11.4, 7.7, 3.6 Hz, 1H), 4.13 (q, *J* = 4.2 Hz, 1H), 3.75 – 3.63 (m, 7H), 3.50 (ddt, *J* = 17.0, 13.5, 6.9 Hz, 4H), 3.37 (dt, *J* = 9.5, 6.2 Hz, 1H), 3.29 – 3.25 (m, 2H), 3.25 – 3.20 (m, 1H), 3.04 (dd, *J* = 10.1, 4.6 Hz, 1H), 2.34 (t, *J* = 6.5 Hz, 2H), 2.28 (ddd, *J* = 13.6, 8.0, 5.5 Hz, 1H), 1.92 (ddd, *J* = 13.7, 3.1, 1.6 Hz, 1H), 1.43 – 1.11 (m, 8H), 1.06 (dd, *J* = 19.3, 6.8 Hz, 12H). **$^{13}$C NMR (176 MHz, $CDCl_3$)** δ 158.37, 157.22, 157.02, 145.04, 136.30, 136.23, 130.17, 128.27, 127.72, 126.65, 117.57, 116.71, 113.04, 104.21, 85.89, 84.40, 84.36, 74.43, 74.33, 67.68, 64.62, 60.41, 58.14, 58.03, 55.22, 40.76, 40.74, 39.83, 29.33, 28.69, 26.34, 25.66, 24.59, 24.55, 24.50, 24.46, 20.34, 20.30. **HRMS**: calculated for $C_{43}H_{57}F_3N_3O_8P$ [M+H]$^+$: 832.38, found: 832.40.

**Scheme S2. Synthesis of ribose-carbon 18 phosphoramidite (R-$C_{18}$)**

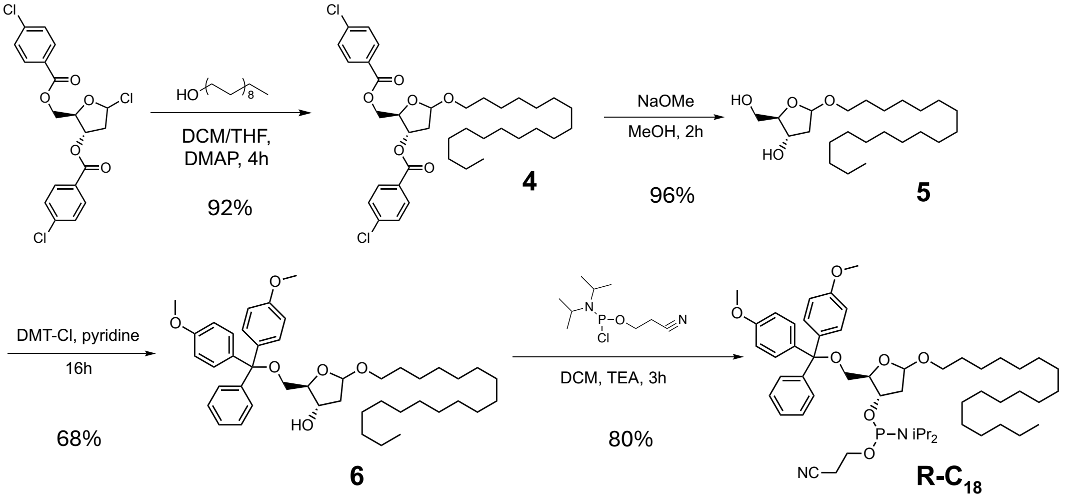


**Synthesis of $C_{18}$-ribose (4, Scheme S2)** 1-Chloro-3,5-di(4-chlorbenzoyl)-2-deoxy-D-ribose (1.00 g, 2.33 mmol, 1 eq.) and DMAP (85.3 mg, 0.70 mmol, 0.3 eq.) was dissolved in 15 mL anhydrous dichloromethane. 1-Octadecanol (0.75 g, 2.80 mmol, 1.2 eq.) was dissolved separately in 5 ml anhydrous tetrahydrofuran and added into the previous mixture. The reaction was allowed to stir at R.T. for at least 4 h before dried under rotary-evaporation. The product was purified by flashing chromatography on silica column with a dichloromethane (0 to 30% ethyl acetate) eluent system. The purified product was dried under high vacuum and yield a white powder. **$^1$H NMR (700 MHz, $CDCl_3$)** δ 7.96 – 7.84 (m, 4H), 7.36 – 7.29 (m, 4H), 5.52 (ddd, *J* = 7.5, 4.7, 2.9 Hz, 1H), 5.26 (dd, *J* = 5.6, 2.4 Hz, 1H), 4.55 – 4.37 (m, 3H), 3.65 (ddt, *J* = 19.9, 9.4, 6.8 Hz, 1H), 3.30 (dt, *J* = 9.6, 6.8 Hz, 1H), 2.47 (ddd, *J* = 14.2, 7.2, 2.4 Hz, 1H), 2.28 (dt, *J* = 14.2, 5.2 Hz, 1H), 1.47 – 1.11 (m, 32H), 0.80 (t, *J* = 7.0 Hz, 3H). **$^{13}$C NMR (176 MHz, $CDCl_3$)** δ 165.44, 165.29, 139.88, 139.60, 131.22, 131.19, 131.12, 131.09, 128.86, 128.80, 128.78, 128.25, 127.92, 104.67, 81.67, 76.03, 68.41, 65.51, 39.19, 39.17, 32.00, 29.80, 29.79, 29.76, 29.72, 29.70, 29.66, 29.60, 29.57, 29.51, 29.47, 26.35, 26.20, 22.79, 14.27. **HRMS**: calculated for $C_{37}H_{52}Cl_2O_6$ $[M+H]^+$: 663.31, found: 663.46.

**Synthesis of deprotected C18-ribose (5, Scheme S2)** Compound **4** (2.44 g, 3.68 mmol, 1 eq.) was dissolved in 18.4 mL 0.5 M NaOMe in methanol solution (9.21 mmol, 2.5 eq.) and stirred at R.T. for 2 h. The reaction mixture was dried by rotary-evaporation and purified by flashing chromatography on silica column with a dichloromethane:ethyl acetate (7:3, v:v) added with methanol (0 to 5%). The purified product was dried under high vacuum and yield a viscous liquid. **$^1$H NMR (700 MHz, $CDCl_3$)** δ 5.21 – 5.14 (m, 1H), 4.13 – 4.07 (m, 1H), 3.67 (dddd, *J* = 15.8, 9.2, 6.6, 4.6 Hz, 2H), 3.57 (ddd, *J* = 32.5, 12.0, 4.2 Hz, 1H), 3.34 (ddt, *J* = 15.8, 9.5, 6.7 Hz, 1H), 2.05 – 1.95 (m, 1H), 1.49 (p, *J* = 5.6 Hz, 2H), 1.18 (d, *J* = 3.3 Hz, 28H), 0.81 (td, *J* = 7.1, 3.7 Hz, 3H). **$^{13}$C NMR (176 MHz, $CDCl_3$)** δ 104.48, 104.45, 104.38, 104.34, 87.77, 87.60, 73.06, 72.36, 68.68, 67.72, 67.70, 63.56, 63.20, 42.89, 41.50, 32.00, 29.79, 29.77, 29.75, 29.70, 29.66, 29.63, 29.46, 26.25, 26.14, 22.79, 14.27. **HRMS**: calculated for $C_{23}H_{46}O_4$ $[M-H]^-$: 385.34, found: 385.33.

**Synthesis of C18-ribose-DMT (6, Scheme S2)** Compound **5** (1.03 g, 2.66 mmol, 1 eq.) was dissolved in 10 mL anhydrous pyridine and cooled with an ice-bath. Dimethoxytrityl chloride (2.02 g, 2.93 mmol, 1.1 eq.) was dissolved in 5 mL anhydrous pyridine and added dropwise into the previous mixture. The reaction was allowed to gradually come back to R.T. and stirred overnight. Pyridine was then removed from the reaction mixture via rotary evaporation (with the addition of toluene), and the mixture was diluted with ethyl acetate. The solution was extracted with 0.1 M $NaHCO_3$ (2x) and brine (2x) and dried with $MgSO_4$ followed by rotary-evaporation. The concentrated solution was then purified by flashing chromatography on silica column with a dichloromethane added with ethyl acetate (0 to 10%) eluent system. The purified product was dried with rotary-evaporation and yield a viscous liquid yellow in color. **$^1$H NMR (700 MHz, $CDCl_3$)** δ 7.38 – 7.10 (m, 9H), 6.77 – 6.71 (m, 4H), 5.05 (dd, *J* = 5.4, 2.1 Hz, 1H), 4.33 (tt, *J* = 6.8, 3.4 Hz, 1H), 3.86 (dt, *J* = 6.9, 5.0 Hz, 1H), 3.70 (s, 6H), 3.50 (dt, *J* = 9.4, 6.8 Hz, 1H), 3.25 – 3.17 (m, 2H), 3.07 (dd, *J* = 9.4, 6.9 Hz, 1H), 2.09 (ddd, *J* = 13.2, 6.8, 2.1 Hz, 1H), 1.94 (ddd, *J* = 12.7, 6.6, 5.4 Hz, 1H), 1.74 (d, *J* = 3.7 Hz, 1H), 1.38 – 1.05 (m, 32H), 0.80 (t, *J* = 7.0 Hz, 3H). **$^{13}$C NMR (176 MHz, $CDCl_3$)** δ 158.48, 144.87, 136.13, 136.10, 130.06, 130.04, 129.15, 128.19, 127.87, 127.83, 127.78, 126.79, 113.12, 103.82, 86.20, 84.53, 73.63, 67.89, 65.28, 55.26, 55.20, 41.19, 31.95, 29.73, 29.70, 29.68, 29.64, 29.62, 29.60, 29.57, 29.48, 29.38, 22.71, 14.14. **HRMS**: calculated for $C_{44}H_{64}O_6$ $[M+H]^+$: 689.47, found: 688.47.

**Synthesis of R-$C_{18}$ (R-$C_{18}$, Scheme S2)** Compound **6** (0.39 g, 0.57 mmol, 1 eq.) and triethylamine (276.1 µL, 1.98 mmol, 3.5 eq.) were dissolved in 8 mL anhydrous dichloromethane and cooled in an ice-bath. 2-Cyanoethyl N,N-diisopropylchlorophosphoramidite (315.6 µL, 1.42 mmol, 2.5 eq.) was dissolved in 2 mL anhydrous dichloromethane and added into the reaction mixture. The reaction was reacted on ice for 30 min and R.T. for 2.5 h. Ethyl acetate was added to dilute the reaction and extracted with 0.1 M $NaHCO_3$ (2x) and brine (2x). The solution was dried with $MgSO_4$ and concentrated via rotary-evaporation. The mixture was purified by flashing chromatography on silica column with a hexane added with ethyl acetate (0 to 20%) eluent system. The final product is dried under high vacuum and yield a viscous liquid. **$^1$H NMR (700 MHz, $CDCl_3$)** δ 7.53 – 7.28 (m, 7H), 7.28 – 7.19 (m, 2H), 6.86 – 6.80 (m, 4H), 5.23 (dd, *J* = 5.2, 2.8 Hz, 1H), 4.46 (dtd, *J* = 10.2, 6.1, 4.2 Hz, 1H), 4.17 (q, *J* = 5.0 Hz, 1H), 3.80 (s, 7H), 3.75 (ddt, *J* = 10.2, 8.0, 6.7 Hz, 1H), 3.68 (dt, *J* = 9.4, 6.9 Hz, 1H), 3.55 (dh, *J* = 10.3, 6.7 Hz, 2H), 3.35 (dt, *J* = 9.4, 6.8 Hz, 1H), 3.21 (dd, *J* = 9.8, 4.9 Hz, 1H), 3.16 (dd, *J* = 9.8, 5.8 Hz, 1H), 2.61 (td, *J* = 6.5, 2.2 Hz, 2H), 2.24 – 2.14 (m, 2H), 1.47 (qt, *J* = 9.5, 4.3 Hz, 2H), 1.35 – 1.29 (m, 3H), 1.28 (s, 14H), 1.25 (d, *J* = 14.3 Hz, 13H), 1.16 (d, *J* = 6.8 Hz, 7H), 1.07 (d, *J* = 6.8 Hz, 6H), 0.90 (t, *J* = 7.0 Hz, 3H). **$^{13}$C NMR (176 MHz, $CDCl_3$)** δ 158.38, 144.99, 136.26, 136.24, 130.15, 130.14, 128.30, 127.71, 126.63, 117.52, 113.01, 104.17, 85.93, 84.40, 84.37, 74.60, 74.50, 68.05, 64.91, 58.24, 58.13, 55.17, 43.20, 43.13, 40.76, 40.74, 31.94, 29.73, 29.68, 29.64, 29.53, 29.38, 26.16, 24.60, 24.56, 24.50, 24.46, 22.71, 20.32, 20.28, 14.14. **HRMS**: calculated for $C_{53}H_{81}N_2O_7P$ $[M+H]^+$: 889.58, found: 889.59.

**Synthesis of polynorbornene bottlebrush (pac-PN)**

Norbornenyl bromide and norbornenyl PEG were synthesized following previously reported procedures.[1] Norbornenyl bromide (5 eq.) was dissolved in deoxygenated DCM under a nitrogen atmosphere. The 2nd generation Grubbs catalyst (1 eq.), dissolved in deoxygenated DCM, was then added to the norbornenyl bromide solution. The mixture was stirred vigorously for 30 minutes, then norbornenyl PEG (50 eq.) in deoxygenated DCM was added. The reaction was stirred for an additional 6 hours. 10 µL of reacted mixture was lyophilized and then dissolved in 0.05 M lithium bromide in HPLC-grade DMF to be analyzed by DMF GPC for molecular weight and PDI measurement. Several drops of ethyl vinyl ether were added to quench the reaction, followed by 2 hours of stirring. The solution was concentrated under vacuum, and the resulting residue was precipitated into cold diethyl ether three times. The precipitate was dried under vacuum, yielding a white powder. The pac-PN brush polymer was then reacted with excess sodium azide in anhydrous DMF overnight at room temperature. The mixture was dialyzed against Nanopure™ water for one week and lyophilized to obtain a white powder. To synthesize pac-PN-ASO, the azide-functionalized PN bottlebrush polymer (50 nmol) was dissolved in 1 mL of 2 M sodium chloride solution and reacted with dibenzocyclooctyne (DBCO)-modified ASO (50 nmol) at 50 °C overnight. The conjugate was purified by aqueous GPC, desalted, and lyophilized. The purified PN-pacDNA was stored at -20 °C until use.

**Oligonucleotide and ribose-based pacDNA backbone synthesis**

All oligonucleotide and ribose backbones were synthesized on a Dr. Oligo 48 DNA synthesizer (Biolytic Lab Performance, Inc., Fremont, CA). The time for the coupling of ribose-phosphoramidites were set at 10 min (compared to 15 s for regular phosphoramidite). Oligonucleotide strands were cleaved from the CPG support and deprotected in aqueous ammonium hydroxide solution (28-30% $NH_3$ basis) at room temperature for 16 h. Free oligonucleotide were purified by RP-HPLC, followed by the removal of dimethoxytrityl (DMT) groups using 20% acetic acid. The oligonucleotide with ribose backbones were purified with Glen-Pak™ DNA purification cartridge (for use with disposable syringes) (Glen Research Co., USA.) following the manufacture's protocol.

**PEGylation of ribose-based pacDNAs**

The purified backbone (50 nmol, 1 eq.) and NHS-terminated 10kDa mPEG (50 nmol, 1 eq.) were dissolved in 800 µL of phosphate buffered saline (PBS, pH 7.4). The mixture was shaken overnight at 4 °C and then desalted using Nap-Column. 50 equivalents of tetrabutylammonium nitrate were added to the desalted product. The desalted product was vacuum dried and then dissolved in 1 mL anhydrous N, N-Dimethylformamide (DMF) containing 42 µL triethylamine. To this mixture, 1 equiv of NHS-terminated PEG (dissolved in 1 mL DMF) was added in 5 aliquots (12 h between each aliquot), and the mixture was gently shaken at room temperature. The mixture was dried *in vacuo* and purified by aqueous GPC.

**Cell culture and animals**

NCI-H358 cells was cultured in RPMI 1640 media supplemented with 10% fetal bovine serum (FBS) and 1% antibiotics. K273 cells were cultured in DMEM media supplemented with 5% fetal bovine serum (FBS), 10µg/mL Epidermal Growth Factor (EGF), 1mg/mL hydrocortisone, 1 mg/mL cholera toxin, 4mg/mL insulin, and 1% penicillin-streptomycin (PS). All cells were cultured at 37 °C in a humidified atmosphere containing 5% $CO_2$.

C57BL/6 mice (6-8 weeks old) were purchased from Charles River Laboratory., USA. Animals were housed at Northeastern University animal facilities. Animal protocols (protocol number: 22-0309R) were approved by the Institutional Animal Care and Use Committee of Northeastern University and carried out in accordance with the approved guidelines.

**Cellular uptake**

NCI-H358 cells were cultured in a 24-well plate with a cell seeding density of $1.0´10^6$ cells per well in 1 mL complete RPMI medium for 24 h at 37 °C. Next, cells were washed 2´ with 1´ PBS and treated with fluorescently labeled test agents and controls, which were dissolved in serum-free RPMI culture medium at various concentrations (1 μM to 5 μM). Cells were further incubated with the samples for 4 h at 37 °C, before being washed with 1´ PBS 3´ and treated with 100 μL of 0.25% trypsin/EDTA solution (Gibco, USA) followed by 700 μL of cold 1´ PBS. Detached cells were collected for flow cytometry analysis (CytoFLEX Flow Cytometer, Beckman Coulter, USA.).

**Confocal microscopy**

NCI-H358 cells were cultured in a 24-well glass bottom plate at $1.0\times10^5$ cells per well in 1 mL complete RPMI medium for 24 h at 37 °C. On the following day, cells were washed with 1´ PBS 3´, and fluorescently labeled test agents and controls dissolved in serum-free RPMI culture medium at were added to give 5 μM of total DNA. Cells were further incubated at 37 °C for 8 h. Next, cells were washed with 1´ PBS 3´ and fixed with 4% paraformaldehyde for 30 min at R.T., followed by another 3´ wash with 1´ PBS. Finally, the cells were stained with DAPI for 10 min before imaged on an LSM-880 confocal laser scanning microscope (Carl Zeiss Ltd., Cambridge, UK). All imaging settings were kept identical in each study.

**Cell viability**

The cell viability of NCI-H358 and K273 after treatment with ribose-based pacDNA and bottlebrush polymer was analyzed by MTT (dimethylthiazol-diphenyltetrazolium bromide) colorimetric assay. Cells were seeded in 96-well plates at a density of $1\times10^4$ cells per well in 175 μL full growth media and cultured for 24 h at 37 °C with 5% $CO_2$. Then cells were treated with pacDNA and bottlebrush polymer in the concentration range of 0.1 – 10 μM (equiv. of DNA). Cells treated with vehicle served as a control. After 48 h of incubation, 20 μL of 5 mg/mL MTT stock solution in PBS was added to each well. After incubation for another 4 h, the media was carefully removed. The resulting blue formazan crystals were dissolved in DMSO (200 μL per well) and measured at 490 nm on a BioTek® Synergy™ Neo2 Multi-Mode microplate reader (BioTek Inc., VT, USA).

**Western blot analysis**

Cells were seeded in 24-well plates at a density of $2.0\times10^5$ cells per well in 1 mL full growth media and cultured for 24 h at 37 °C with 5% $CO_2$. After washing by PBS 1×, pacDNA and bottlebrush polymer (1 μM – 10 μM equiv. of DNA) dissolved in full media (1 mL) was added, and cells were further incubated at 37 °C for 72 h. Next, cells were harvested and whole cell lysates were collected in 100 μL of RIPA cell lysis buffer supplemented with 1% phosphate inhibitor and 1% phosphatase inhibitor. Total proteins in cell lysate were quantified using a bicinchoninic acid (BCA) protein assay kit. Equal amounts of total proteins were separated on a 10% SDS-PAGE gel and electro-transferred to nitrocellulose membrane. The membrane was then blocked with 5% bovine serum albumin (BSA) in Tris-buffered saline supplemented with 0.05% Tween-20 (TBST). After blocking, the membrane was incubated with appropriate primary antibodies overnight at 4 °C. After washing with TBST for three times (10 min per time), the membrane was incubated with secondary antibodies at room temperature for 1 h. The detected proteins were visualized by chemiluminescence using the ECL Western Blotting Substrate (Bio-rad, MA, USA). Antibodies used in this study were: KRAS antibody (cat. sc-30; Santa Cruz), β-actin (cat. AM4302, Cell Signaling Technology), anti-mouse IgG, HRP-linked antibody (cat. 7076S, Cell Signaling Technology).

**Plasma pharmacokinetics (PK) studies**

Immunocompetent C57BL/6 mice were used to examine the plasma PK of free ASO, pacDNAs. Mice were randomly divided into seven groups (n = 5). Cy5-labeled samples were intravenously (i.v.) administrated via the tail vein at equal ASO dosage (0.5 μmol/kg). Blood samples (25 μL) were collected from the submandibular vein at varying time points (30 min, 2 h, 4 h, 10 h, 24 h, 48 h and 72 h) using BD Vacutainer™ blood collection tubes with lithium heparin. Heparinized plasma was obtained by centrifugation at 3000 rpm for 20 min, aliquoted into a 96-well plate, and measured for fluorescence intensity on a Synergy™ Neo2 Multi-Mode microplate reader (BioTek Instruments Inc., VT, USA). The amounts of ASO in the blood samples were estimated using standard curves established for each sample. To establish the standard curves, samples of known quantities were incubated with freshly collected plasma for 1 h at room temperature before fluorescence was measured.

**Allograft tumor model**

To establish allograft tumor model, approximately $5\times10^5$ K273 cells in 100 μL phosphate buffered saline (PBS) were implanted subcutaneously on the right flank of 6-week-old C57BL/6 (female, n = 5). Mice were monitored for tumor growth and body weight every other day. Once tumor volume reached $100mm^3$, mice were divided into groups to keep initial average tumor volumes of each group very close. The mice received following treatment via tail vein injection: 1) PBS, 2) pac-4×1$C_{18}$-ASO2 (0.5 μmol/kg), 3) pac-4×1$C_{18}$-ASO2 (0.5 μmol/kg), 4) Free ASO2 (0.5 μmol/kg). Samples were injected every four days for four doses. Once the tumor volume reached $1000mm^3$, the mice were euthanized by $CO_2$. Major organs and tumors were collected into 10% neutral buffered formalin for overnight fix and then for histological analysis carried by iHisto Inc.

**Anti-PEG immune response**

Healthy C57BL/6 mice were administrated PBS, pac-4×1$C_{18}$-ASO2, free ASO2 or PEG-KLH every four days for four doses. 50-100µL plasma were collected one week after the last dose. The concentration of anti-PEG IgM/IgG were assessed by ELISA kit (Cat # PEGG-1, Cat #PEGM-1, Life Diagnotics, Inc.) according to the manufacturer's protocol.

**Multiplex analysis of cytokines**

The multiplexing analysis was performed using the Luminex™ 200 system (Luminex, Austin, TX, USA) by Eve Technologies Corp. (Calgary, Alberta). Thirty-two markers were simultaneously measured in the samples using Eve Technologies' Mouse Cytokine 32-Plex Discovery Assay® (MilliporeSigma, Burlington, Massachusetts, USA) according to the manufacturer's protocol. The 32-plex consisted of Eotaxin, G-CSF, GM-CSF, IFNγ, IL-1α, IL-1β, IL-2, IL-3, IL-4, IL-5, IL-6, IL-7, IL-9, IL-10, IL-12(p40), IL-12(p70), IL-13, IL-15, IL-17, IP-10, KC, LIF, LIX, MCP-1, M-CSF, MIG, MIP-1α, MIP-1β, MIP-2, RANTES, TNFα, and VEGF. Assay sensitivities of these markers range from 0.3 – 30.6 pg/mL for the 32-plex. Individual analyte sensitivity values are available in the MilliporeSigma MILLIPLEX® MAP protocol.

**Whole-animal and *ex vivo* organ imaging**

Tumor-bearing C57BL/6 mice were i.v. injected with Cy5-labeled samples at an ASO dose of 0.5 µmol/kg. Then mice were scanned at 1, 4, 8, 24 h, 48 h, and 72 h using an IVIS Lumina II imaging system (Caliper Life Sciences, Inc. MA, USA). To evaluate the biodistribution of pacDNAs and the bottlebrush polymer, mice were euthanized using $CO_2$, and major organs, tissues and the tumor were dissected and imaged using IVIS.

**Biodistribution**

The major organs, tissues and tumor were diced into small piece and weighted. Then, the flesh was homogenized in tissue protein extraction reagent (Thermo Fisher, USA), which is supported with 0.5% Triton X-100, using a BeadBlaster D2400-R refrigerated homogenizer (Benchmark scientific, NJ, USA.). After centrifugation, the supernatants were aliquoted into a 96-well plate and measured for fluorescence intensity on a Synergy™ Neo2 Multi-Mode microplate reader (BioTek Instruments Inc., VT, USA). The amounts of ASO in the supernatant were estimated using standard curves established for each sample.

**All-atom molecular dynamics (MD) simulation**

All-atom molecular dynamics (MD) simulations of pacDNAs were conducted to understand the molecular mechanism of how the lipids distribution affects the energy state of pacDNAs. The four initial pacDNAs with six $C_{18}$ units arranged in different distributions were built using ChemDraw. For these four pacDNA models, all PEG side chains were truncated to a 20 mer. Subsequently, each pacDNA model underwent a refinement process in Maestro, during which hydrogen atoms were added to the structures.[2] The OPLS4 forcefield was used to model pacDNAs.[3] To mimic the aqueous environment, each model was solvated in a rectangular water box using the SPC water model with a minimum water shell of 10 Å from any box edge to the nearest atom of the structure. Sodium (Na+) and chloride (Cl-) ions were placed to neutralize the system. Furthermore, the salt (NaCl) concentration was set to 0.15 M to mimic physiological conditions.

Prior to the production run, each pacDNA structure underwent an extensive minimization and equilibration process, detailed as follows. Each system was first relaxed using the default relaxation protocol from Maestro. After that, the system went through an annealing process by first heating the system from 10 K to 400 K and then cooling to 300 K. Following annealing, each pacDNA model was re-solvated with a refreshed water box to accurately represent the aqueous environment. The updated simulation system was used to conduct a production run for 100 ns at a constant temperature of 300 K with 1 bar pressure (NPT ensemble). The Nosé–Hoover algorithm was used for temperature control, and the Martyna–Tobias–Klein algorithm was used to control the pressure. All molecular dynamics simulations were performed using the GPU-accelerated Desmond engine in Schrodinger molecular modeling suite.[4-6] The simulation trajectories from 90 ns to 100 ns were used to calculate the potential energy for the corresponding pacDNA structure.

A full-length and full-atomistic pacDNA model with the largest spacing (pac-4×1$C_{18}$-ASO1) was built to investigate the conformational characteristics. The initial structure of pacDNA-4×1$C_{18}$ with truncated PEG chain was built using ChemDraw. We extended each PEG chain to 226 mer using VMD and Maestro.[2, 7] After that, we followed the same process as described above to minimize and equilibrate the structure. The same annealing protocol was used to optimize the structure. The simulation system with an optimized structure (~1.6 million atoms) was used to run the production run for 60 ns in an NPT ensemble (300 K, 1 bar).

**Table S1. All ribose-based pacDNA structures and sequences used in this study.**

| Sample ID | Sequence |
|---|---|
| Free ASO1 | 5′-GCT ATT AGG AGT CTT T-3′ |
| pac-0$C_{18}$-ASO1 | 5′-GCT ATT AGG AGT CTT TNN NNN NNN NNN NNN NNN NNN NNN NNN NNN NT-3′ |
| pac-2×1$C_{18}$-ASO1 | 5′-GCT ATT AGG AGT CTT TNN NNN NNN NNS NNN NNN NNN NSN NNN NNN NNN T-3′ |
| pac-4×1$C_{18}$-ASO1 | 5′-GCT ATT AGG AGT CTT TNN NNN NSN NNN NNS NNN NNN SNN NNN NSN NNN NNT-3′ |
| pac-6×1$C_{18}$-ASO1 | 5′-GCT ATT AGG AGT CTT TNN NNN SNN NNS NNN NSN NNN SNN NNS NNN NSN NNN NT-3′ |
| pac-8×1$C_{18}$-ASO1 | 5′-GCT ATT AGG AGT CTT TNN NNS NNN NSN NNS NNN SNN NSN NNS NNN SNN NSN NNN T-3′ |
| pac-10×1$C_{18}$-ASO1 | 5′-GCT ATT AGG AGT CTT TNN SNN SNN NSN NNS NNN SNN NSN NNS NNN SNN NSN NNS NNT-3′ |
| Cy5-ASO1 | 5′-Cy5-GCT ATT AGG AGT CTT T-3′ |
| Cy5-pac-0$C_{18}$-ASO1 | 5′-Cy5-GCT ATT AGG AGT CTT TNN NNN NNN NNN NNN NNN NNN NNN NNN NNN NT-3′ |
| Cy5-pac-2×1$C_{18}$-ASO1 | 5′-Cy5-GCT ATT AGG AGT CTT TNN NNN NNN NNS NNN NNN NNN NSN NNN NNN NNNT-3′ |
| Cy5-pac-4×1$C_{18}$-ASO1 | 5′-Cy5-GCT ATT AGG AGT CTT TNN NNN NSN NNN NNS NNN NNN SNN NNN NSN NNN NNT-3′ |
| Cy5-pac-6×1$C_{18}$-ASO1 | 5′-Cy5-GCT ATT AGG AGT CTT TNN NNN SNN NNS NNN NSN NNN SNN NNS NNN NSN NNN NT-3′ |
| Cy5-pac-8×1$C_{18}$-ASO1 | 5′-Cy5-GCT ATT AGG AGT CTT TNN NNS NNN NSN NNS NNN SNN NSN NNS NNN SNN NSN NNN T-3′ |
| Cy5-pac-10×1$C_{18}$-ASO1 | 5′-Cy5-GCT ATT AGG AGT CTT TNN SNN SNN NSN NNS NNN SNN NSN NNS NNN SNN NSN NNS NNT-3′ |
| pac-3×2$C_{18}$-ASO1 | 5′-Cy5-GCT ATT AGG AGT CTT TNN NNN NNN SSN NNN NNN SSN NNN NNN SSN NNN NNN NT-3′ |
| pac-2×3$C_{18}$-ASO1 | 5′-Cy5-GCT ATT AGG AGT CTT TNN NNN NNN NNS SSN NNN NNN NNN SSS NNN NNN NNN NT-3′ |
| pac-1×6$C_{18}$-ASO1 | 5′-Cy5-GCT ATT AGG AGT CTT TNN NNN NNN NNN NNN NSS SSS SNN NNN NNN NNN NNN NT-3′ |
| Free ASO2 | 5′-CAT GTA AAT ATA GCC CT-3′ |
| pac-4×1$C_{18}$-ASO2 | 5′-CAT GTA AAT ATA GCC CTN NNN NNS NNN NNN SNN NNN NSN NNN NNS NNN NNN T-3′ |
| pac-4×1$C_{18}$-scramble | 5′-ACA ACG CTG CAT TAT ATN NNN NNS NNN NNN SNN NNN NSN NNN NNS NNN NNN T-3′ |

N: R-$NH_2$ phosphoramidite (each PEGylated with 10k PEG); S: R-$C_{18}$ phosphoramidite

**Table S2. DMF-GPC analyses for the ribose-based pacDNAs**

| Sample ID | $M_n$(kDa) | $M_w$(kDa) | PDI |
|---|---|---|---|
| pac-**0**$C_{18}$-ASO1 | 362.5 | 438.6 | 1.21 |
| pac-**2×1**$C_{18}$-ASO1 | 377.1 | 467.6 | 1.24 |
| pac-**4×1**$C_{18}$-ASO1 | 379.9 | 474.9 | 1.25 |
| pac-**6×1**$C_{18}$-ASO1 | 379.3 | 466.5 | 1.23 |
| pac-**8×1**$C_{18}$-ASO1 | 391.9 | 474.2 | 1.21 |
| pac-**10×1**$C_{18}$-ASO1 | 394.1 | 492.6 | 1.25 |
| pac-**3×2**$C_{18}$-ASO1 | 387.7 | 473.0 | 1.22 |
| pac-**2×3**$C_{18}$-ASO1 | 385.6 | 478.1 | 1.24 |
| pac-**1×6**$C_{18}$-ASO1 | 388.8 | 478.2 | 1.23 |

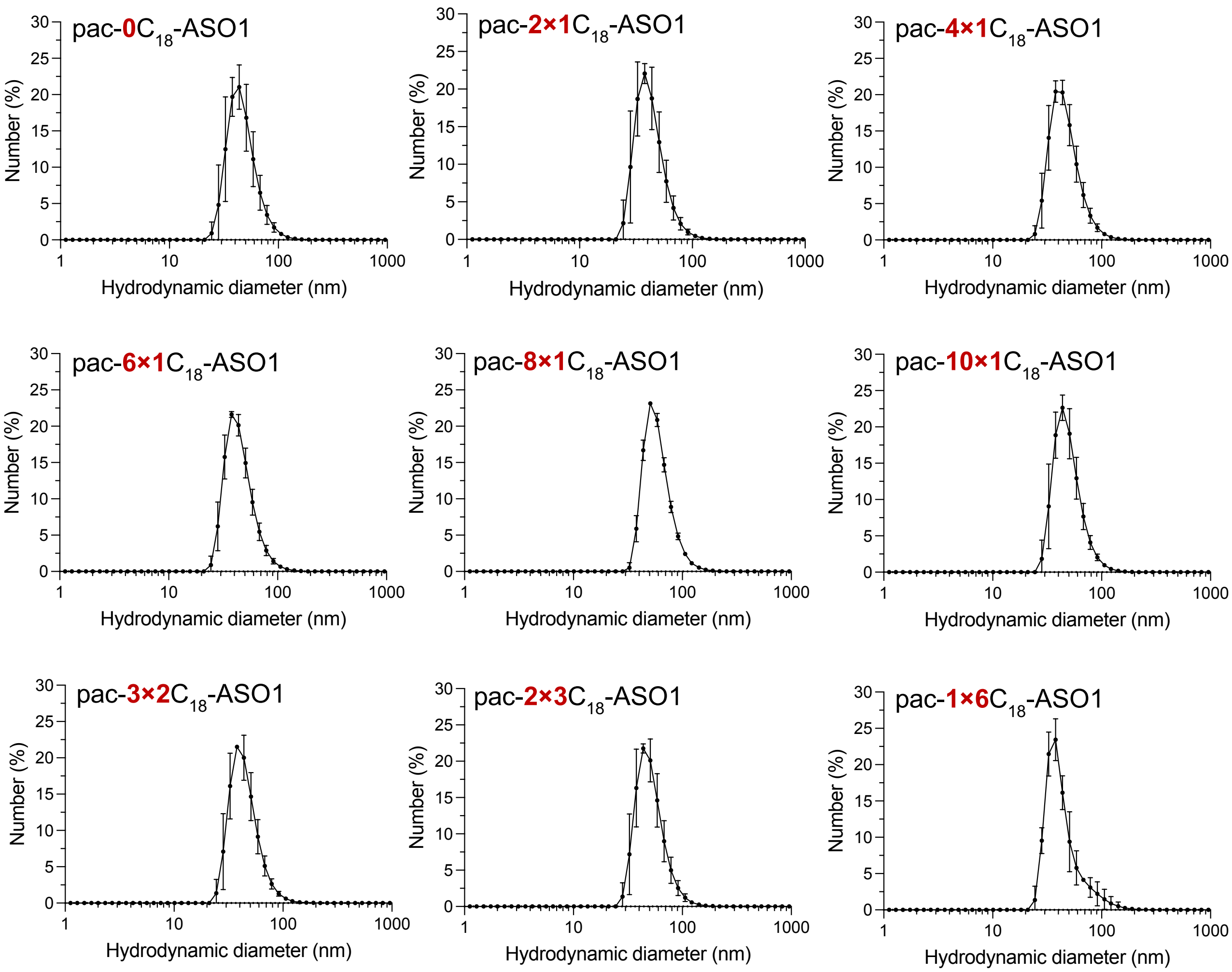


**Figure S1.** DLS measurements of ribose-based pacDNAs in Nanopure™ water.

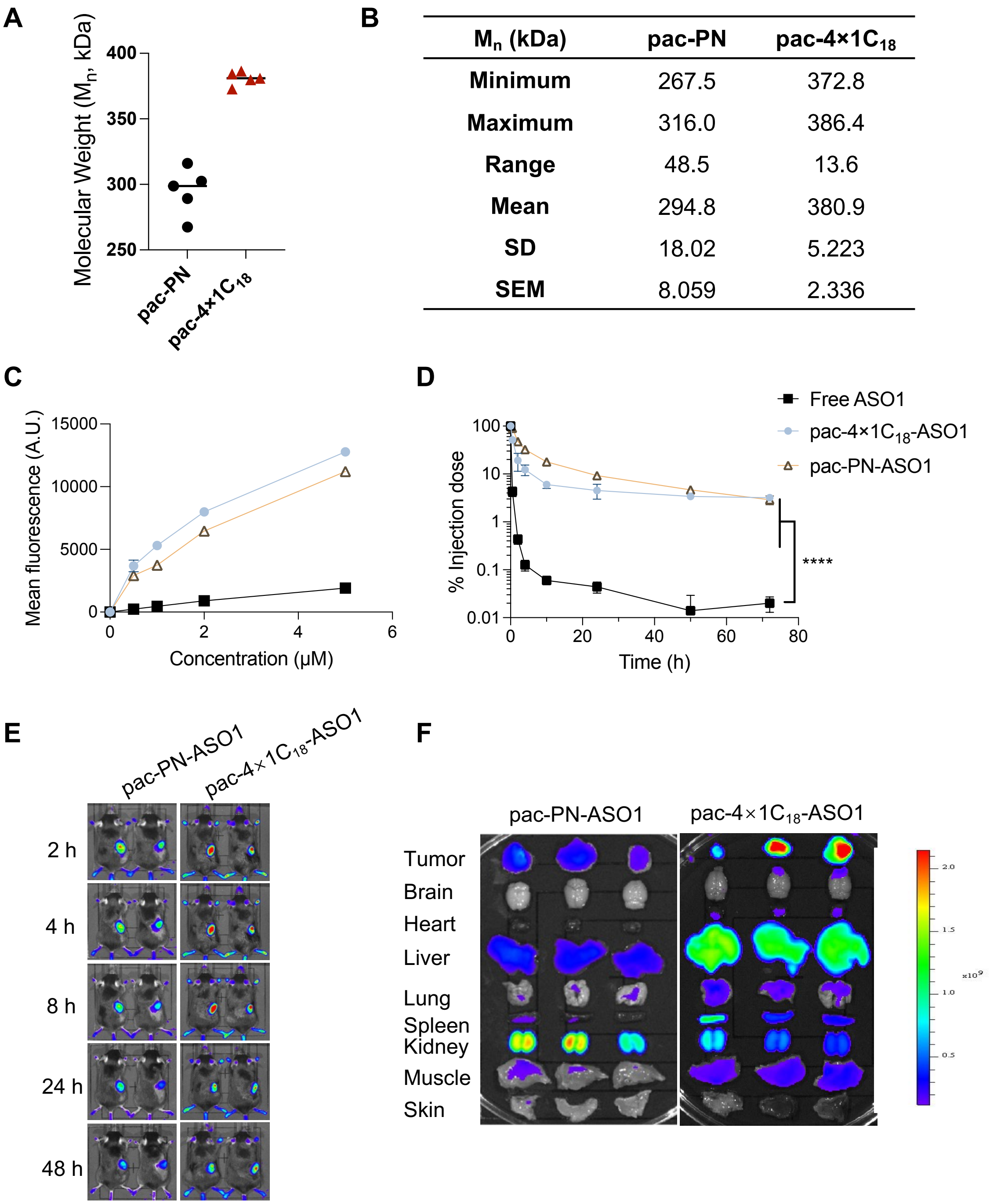


**Figure S2**. The comparison of the polynorbornene bottlebrush (pac-PN) and ribose-based pac $4{\times}1C_{18}$. (A) The molecular weight of five batches of pac-PN and pac $4{\times}1C_{18}$ brush polymers. (B) Descriptive statistics of the molecular weight of (A). (C) Cellualr uptake by NCI-H358 cells of Cy5-labeled pacDNAs and free oligo. (D) Plasma pharmacokinetics of Cy5-labeled pacDNA and free DNA in C57BL/6 mice following i.v. injection. (E) Live animal fluorescence imaging of C57BL/6 mice bearing K273 allograft following i.v. injection of Cy5-labeled pacDNA. Areas surrounding the allograft has been shaved to facilitate imaging. (F) The *ex vivo* imaging of pacDNAs in major organs/tissues of tumor-bearing C57BL/6 mice 72 h post i.v. injection.

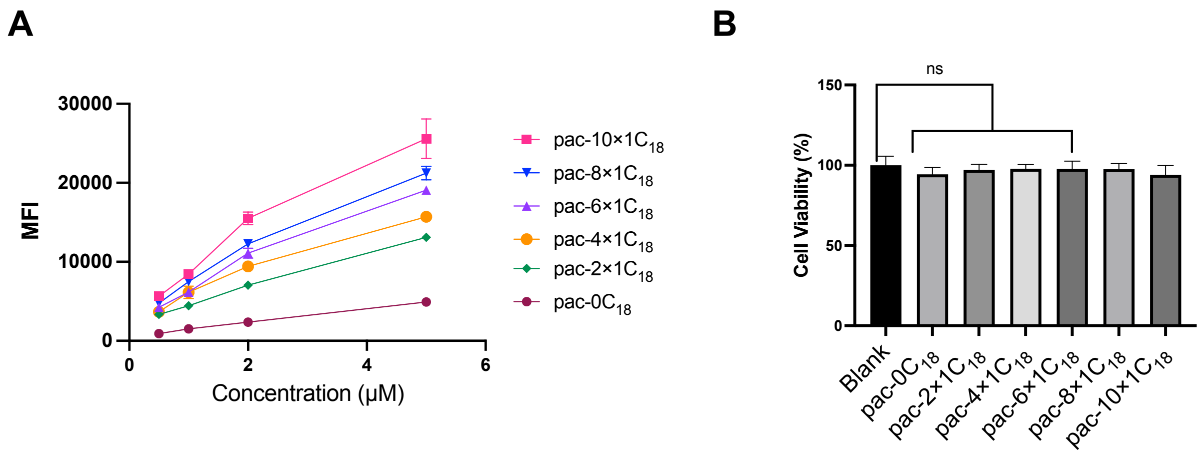


Figure S3. *In vitro* properties of ribose-based pacDNAs without ASO. (A) Cellular uptake by NCI-H358 cells of Cy5-labeled pacDNAs containing varying numbers and arrangement of $C_{18}$ modifiers after 4 h incubation, as determined by flow cytometry. MFI: mean fluorescence intensity. (B) Cytotoxicity of 10μM ribose-based pacDNAs (without ASO) on the proliferation of NCI-H358 cells after 48 h incubation.

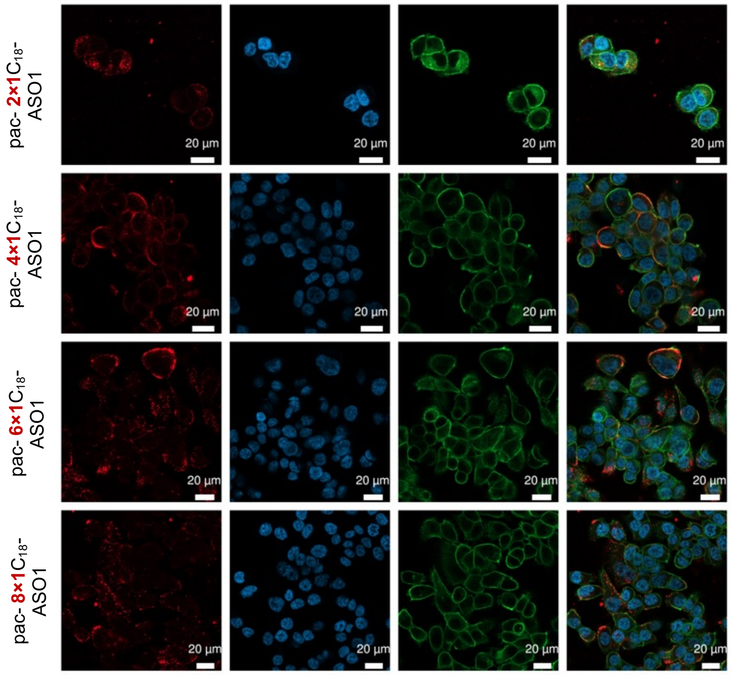


**Figure S4**. Confocal microscopy of NCI-H358 cells treated with Cy5-labeled free DNA or ribose-based pacDNAs with four or ten $C_{18}$ modifiers for 8 h.

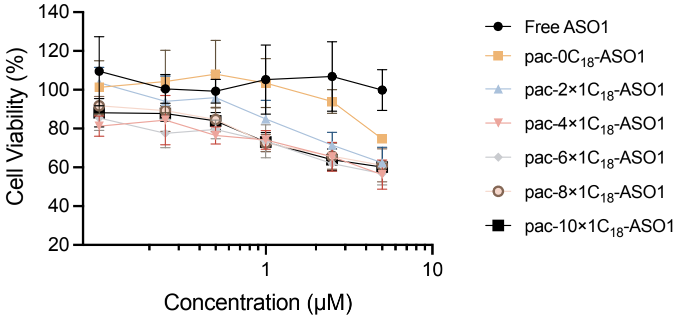


**Figure S5**. Cell viability inhibition efficiency of ribose-based pacDNAs at different concentration (from 100nM to 5μM).

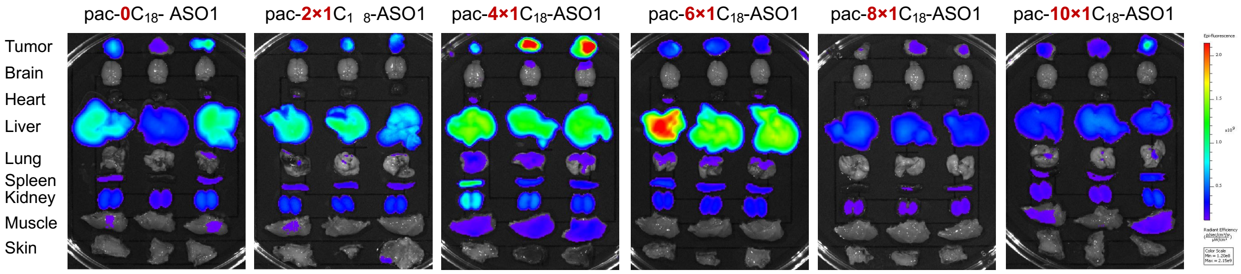


**Figure S6**. Fluorescence imaging of major organs/tissues of tumor-bearing C57BL/6 mice 72h post i.v. injection of Cy5-label ribose-based pacDNAs (with variant number of $C_{18}$ modified on backbone).

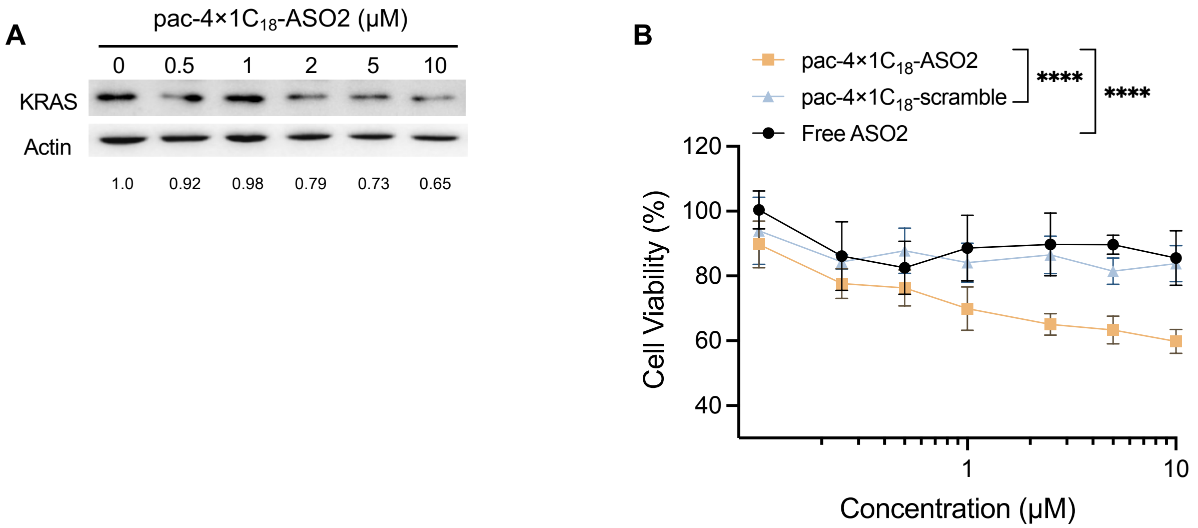


**Figure S7**. In vitro efficacy test of ribose-based pacDNAs on K273 cell line. (**A**) KRAS depletion efficacy of pac-4×1$C_{18}$-ASO2 dose-dependently. (**B**) Inhibition of K273 cell proliferation 48h after treatment of pac-4×1$C_{18}$-ASO2, pac-4×1$C_{18}$-scramble and free-ASO2 at different levels of dose. Statistical analysis was performed using two-way ANOVA. **** $P$ <0.0001.

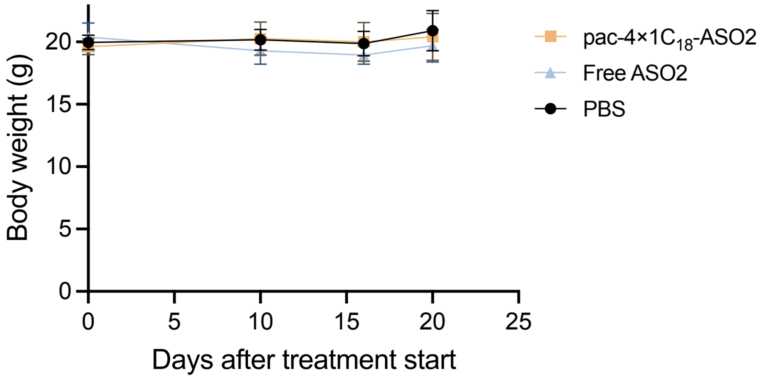


**Figure S8**. Body weight change of C57BL/6 mice bearing K273 allograft model. See figure 5A for the administration frame.

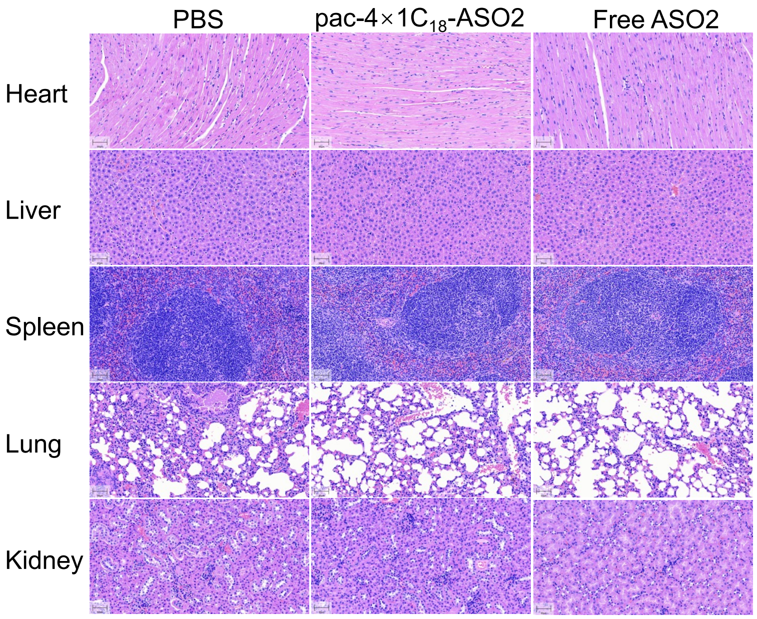


**Figure S9.** H&E staining of major organs and tissues of C57BL/6 mice after i.v. injection of PBS, free ASO2 and pac-4×1$C_{18}$-ASO2 at doses 0.5 μmol/kg every four days, four doses at total.

#### Compound 1

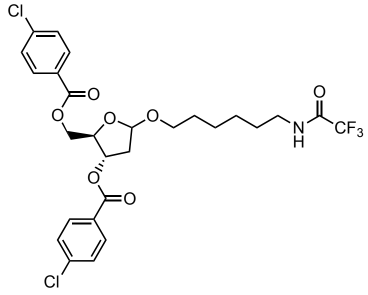

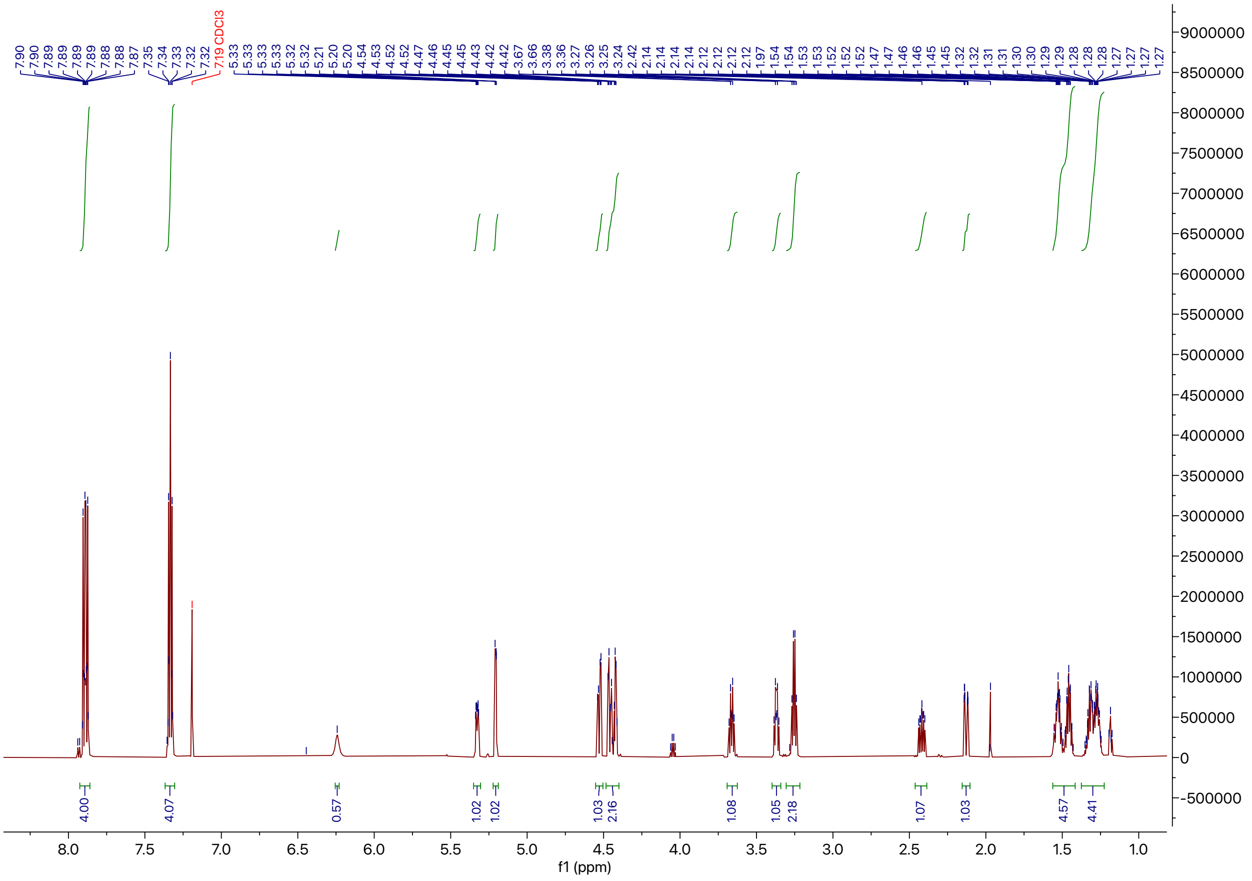

**Figure S10.** $^{1}$H NMR spectrum of compound 1.

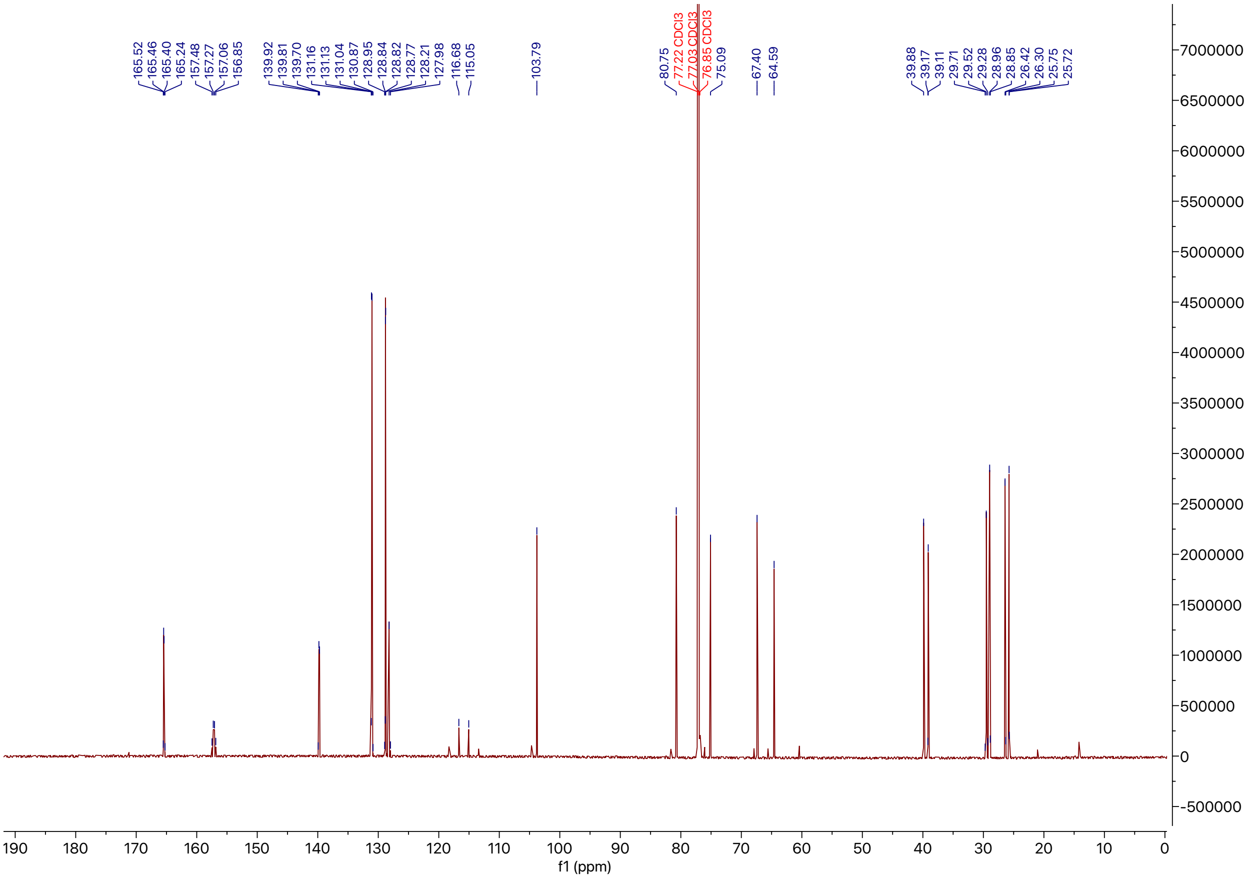

**Figure S11.** $^{13}$C NMR spectrum of compound 1.

#### Compound 2

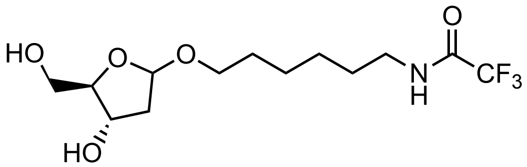

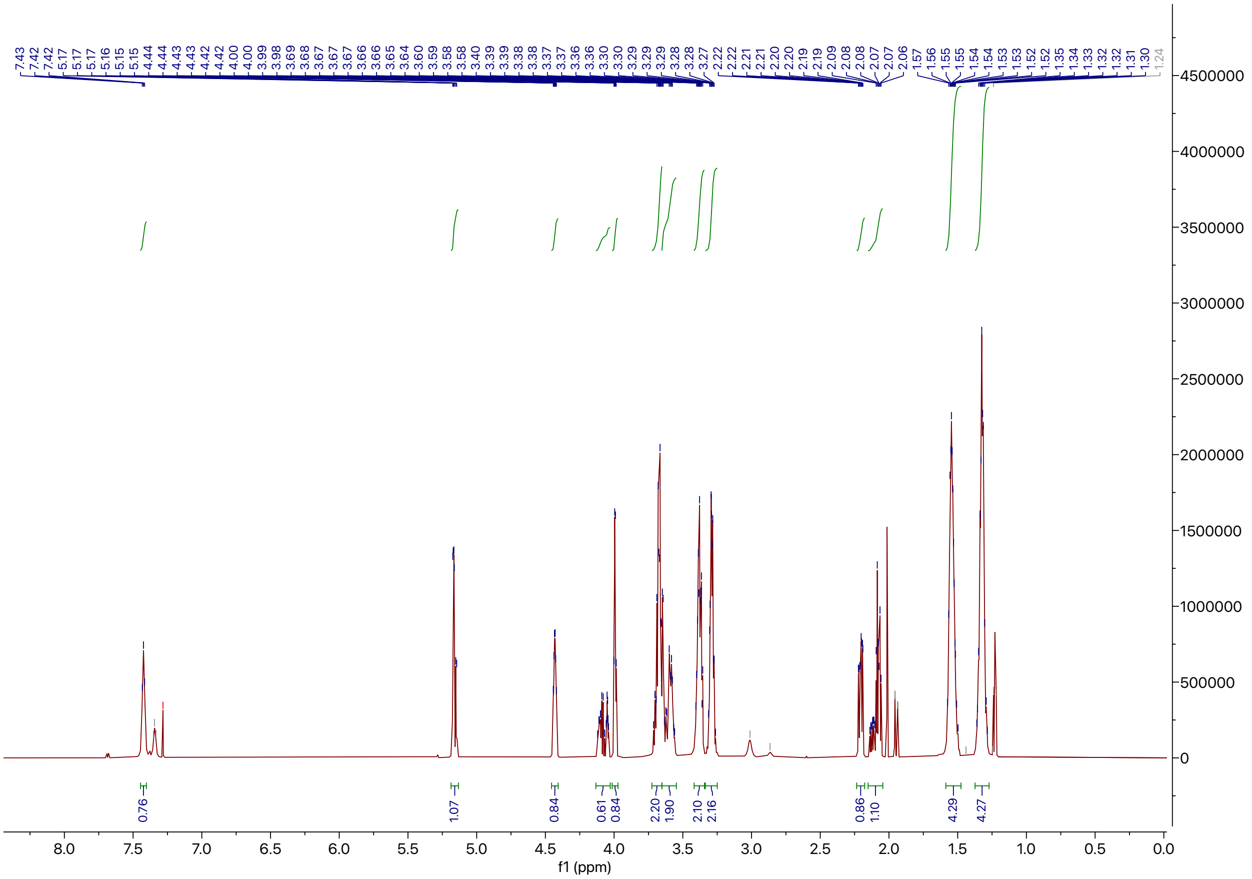

**Figure S12.** $^{1}H$ NMR spectrum of compound 2.

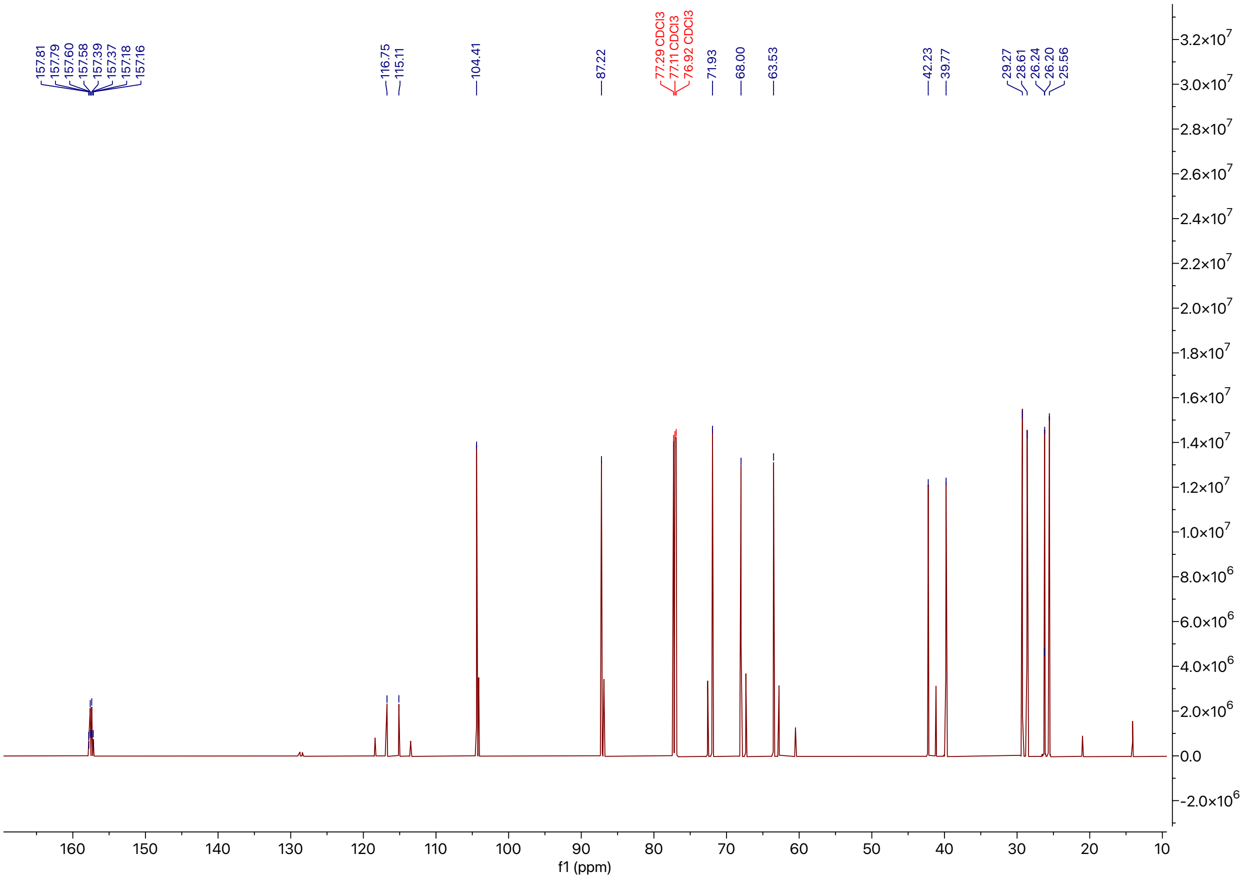


**Figure S13.** $^{13}C$ NMR spectrum of compound 2.

### Compound 3

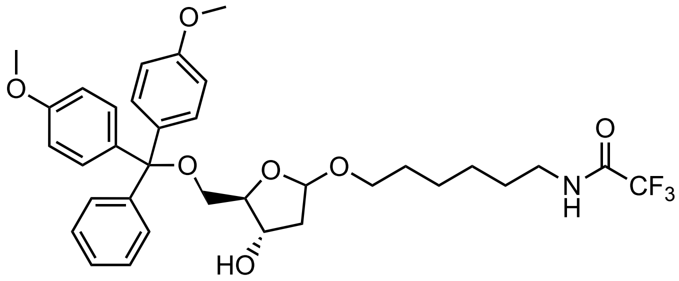


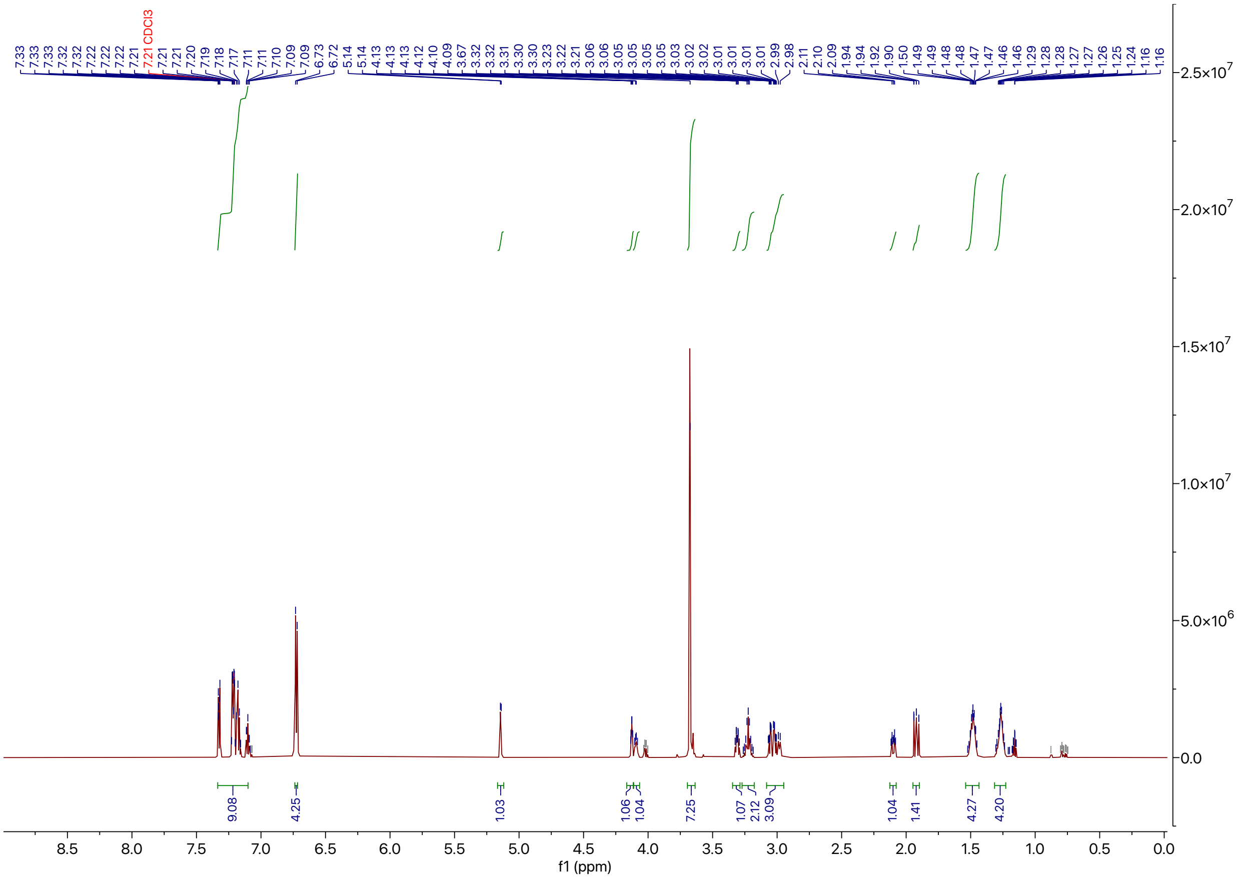


**Figure S14.** $^{1}$H NMR spectrum of compound 3.

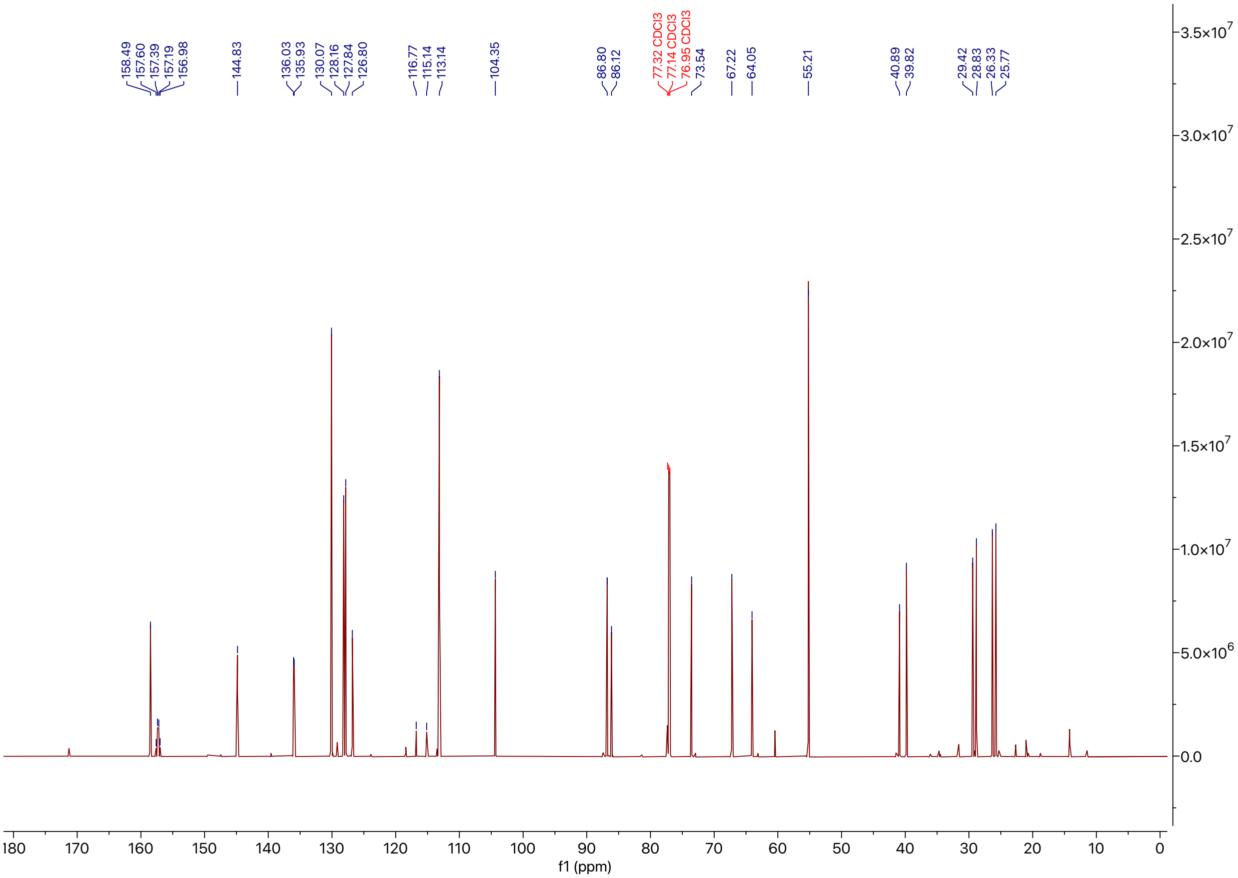


**Figure S15.** $^{13}C$ NMR spectrum of compound 3.

**R-NH$_2$**

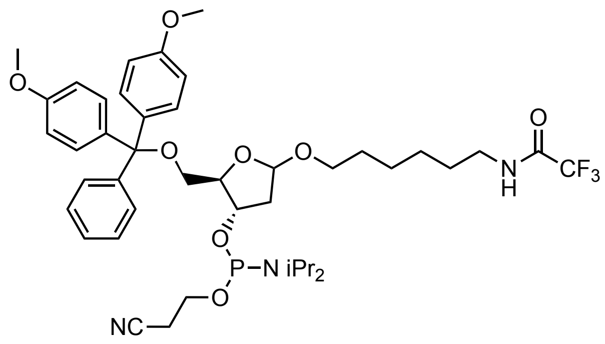

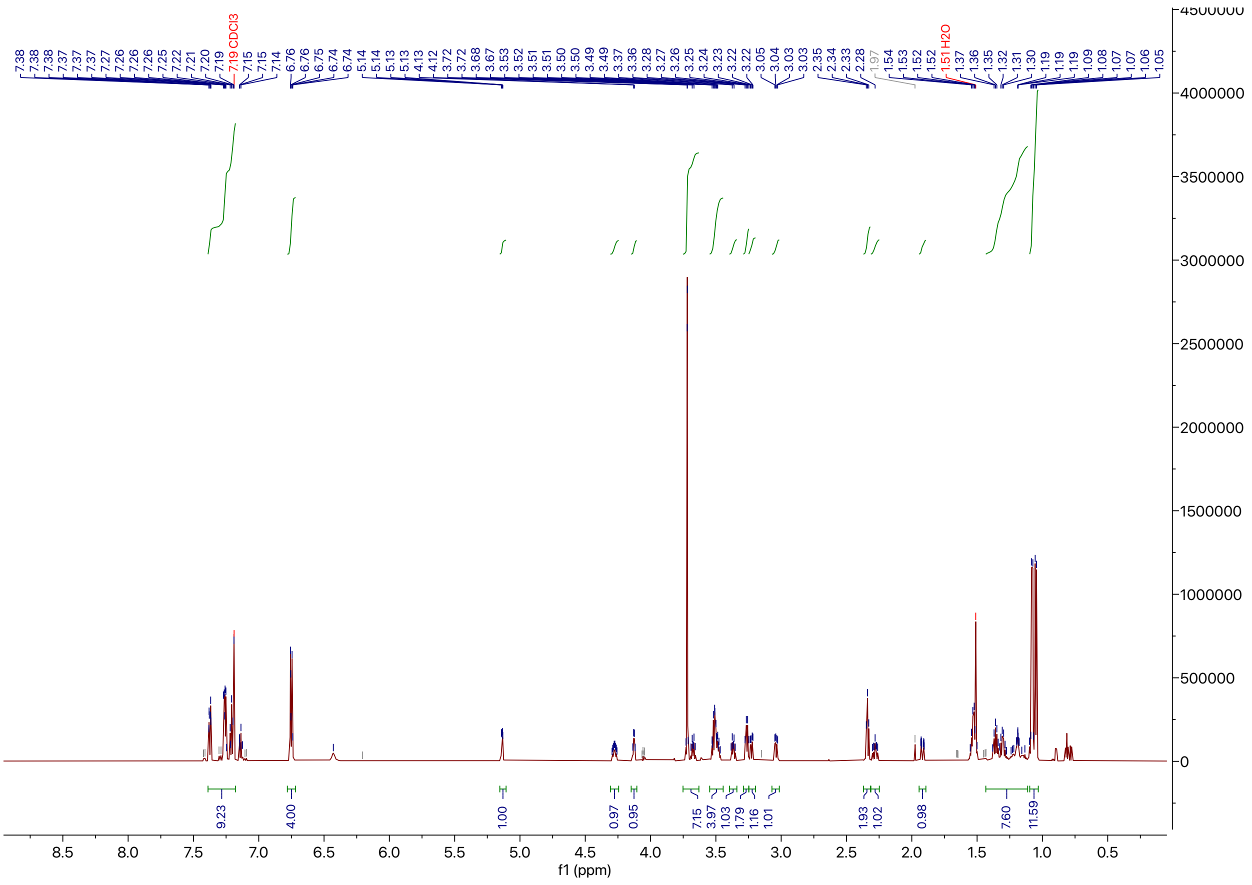

**Figure S16.** $^{1}H$ NMR spectrum of R-NH$_2$.

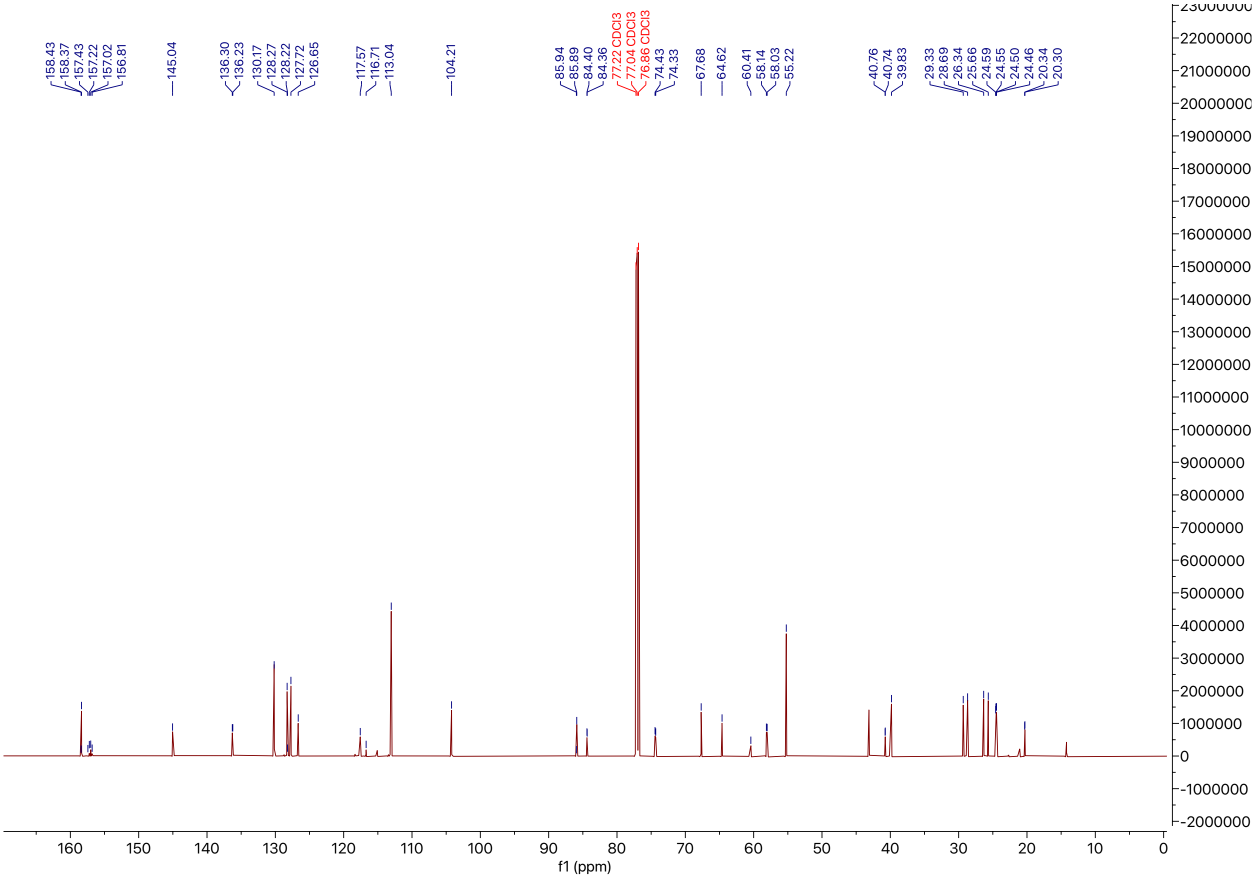


**Figure S17.** $^{13}$C NMR spectrum of R-$NH_2$.

### Compound 4

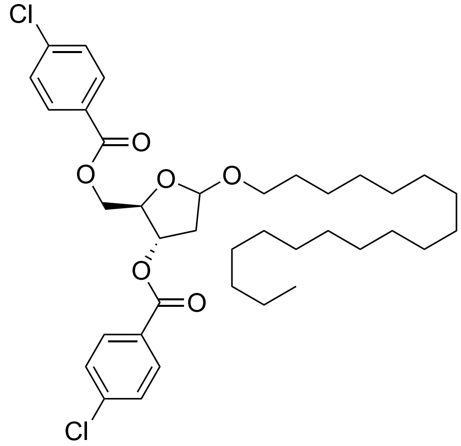


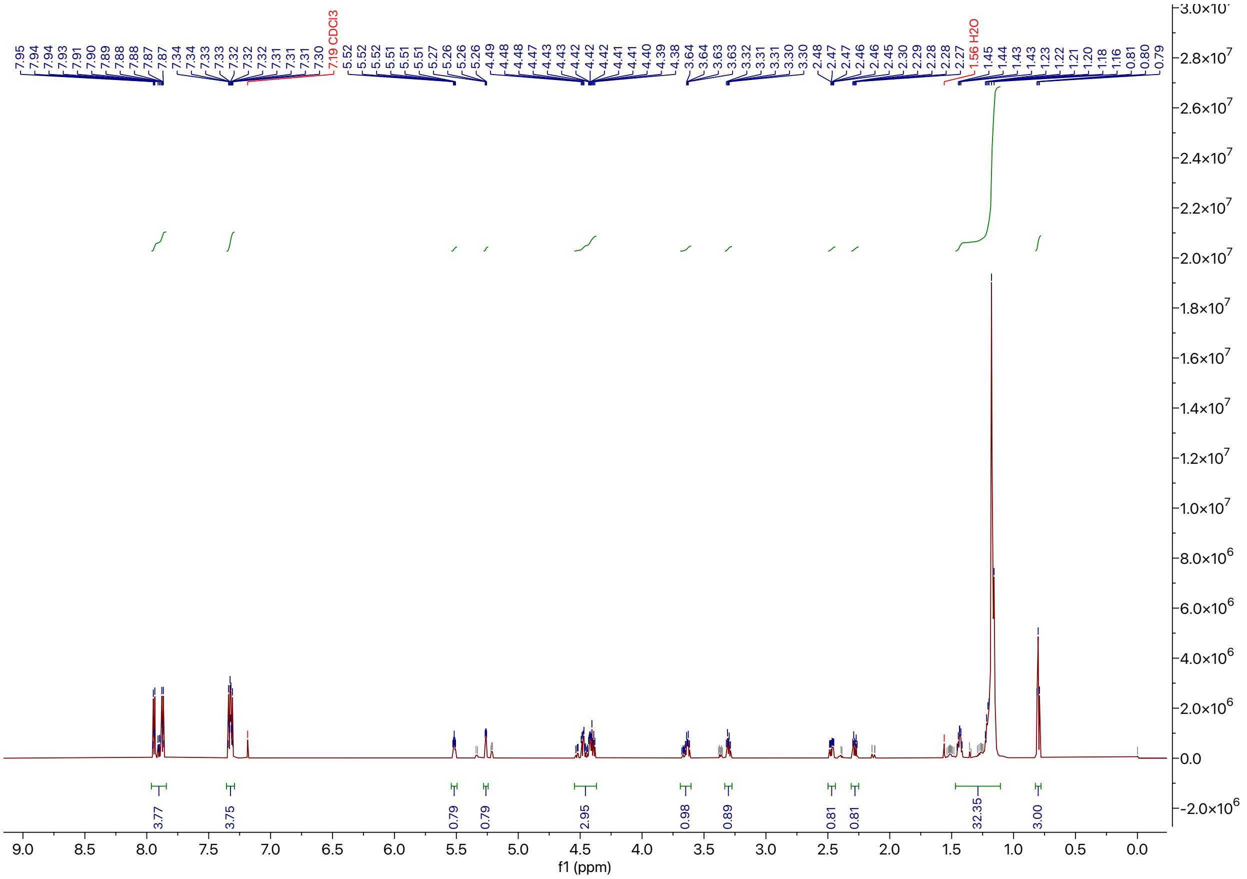


**Figure S18.** $^{1}H$ NMR spectrum of compound 4.

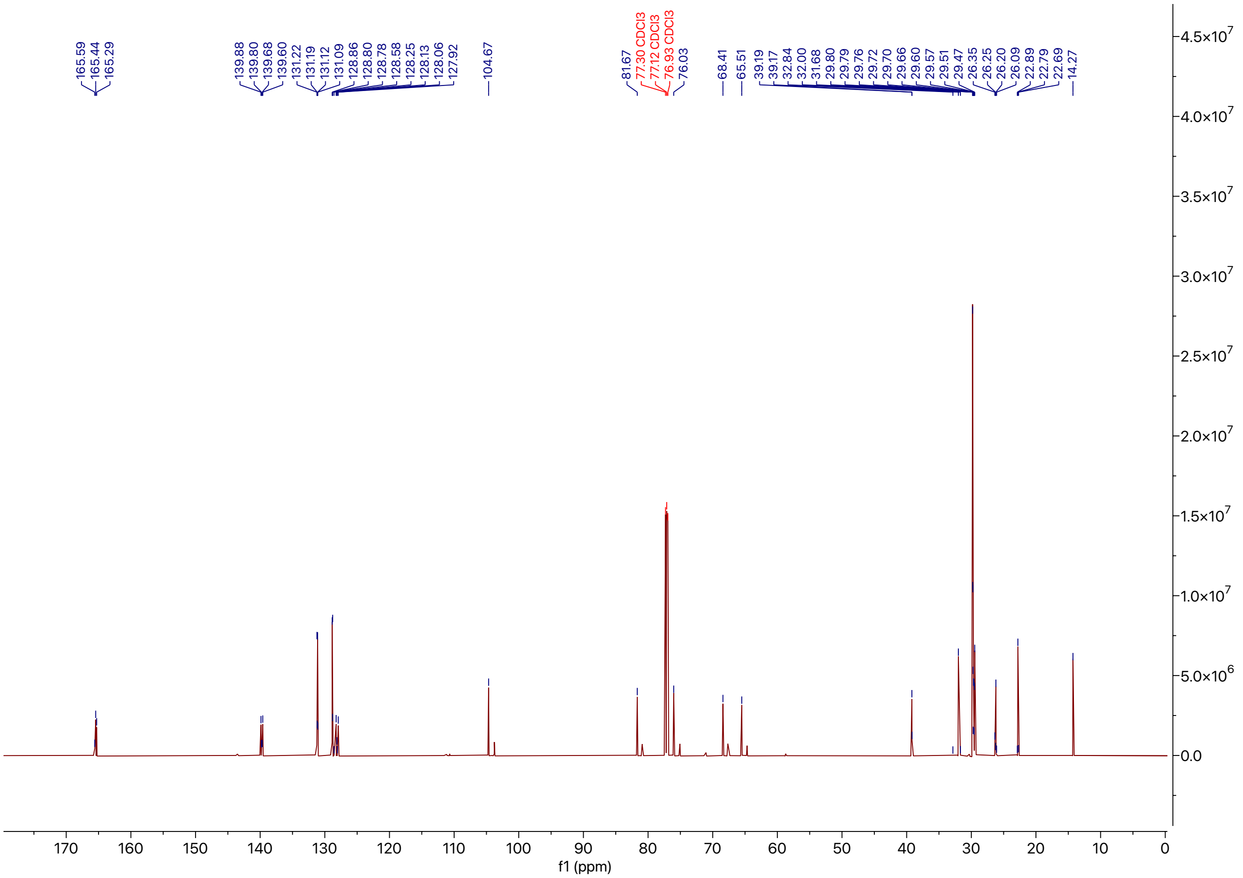


**Figure S19.** $^{13}$C NMR spectrum of compound 4.

### Compound 5

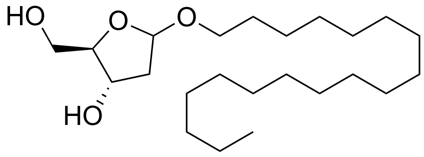


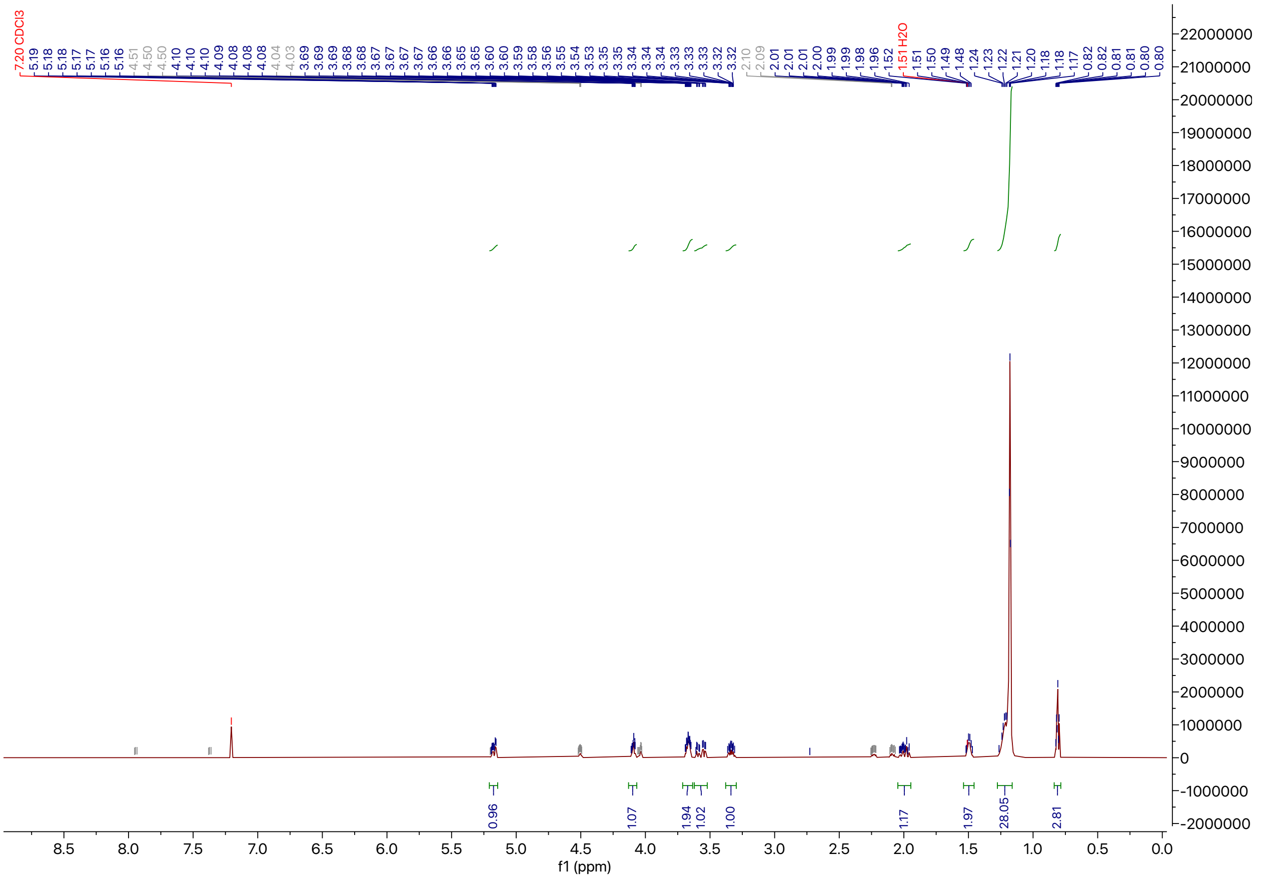


**Figure S20.** $^{1}$H NMR spectrum of compound 5.

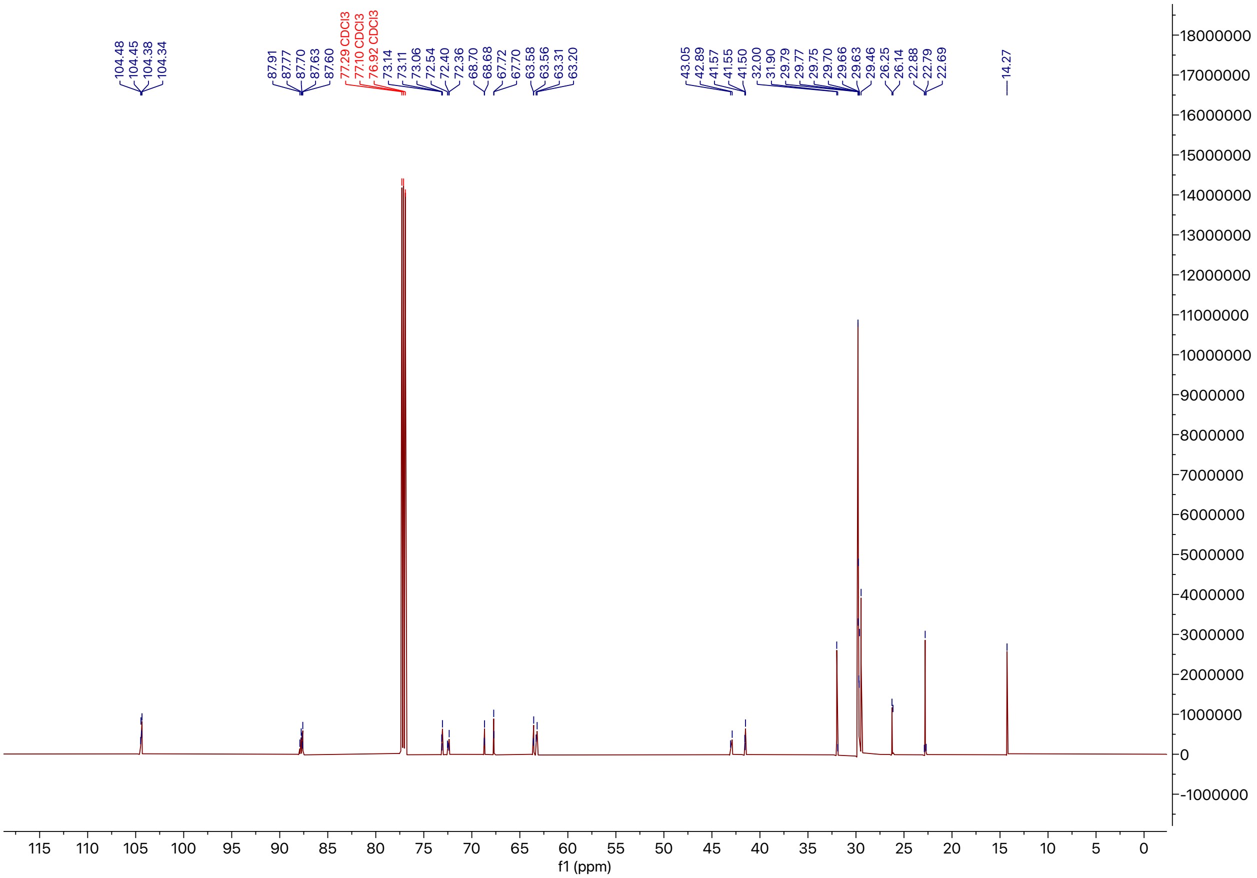

**Figure S21.** $^{13}$C NMR spectrum of compound 5.

### Compound 6

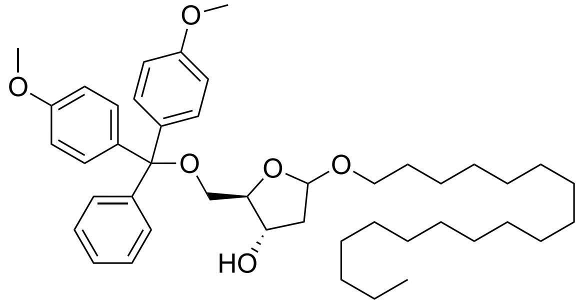

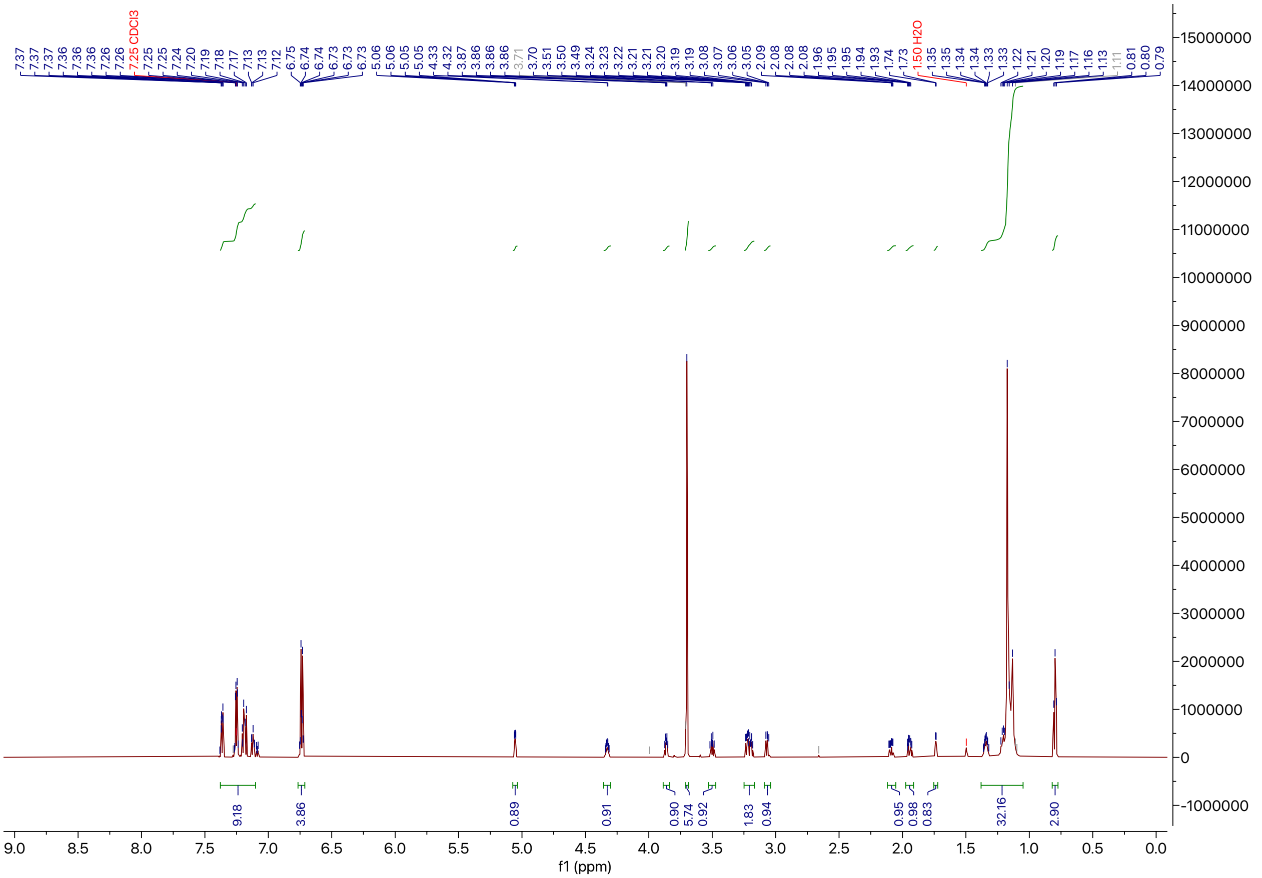


**Figure S22.** $^1$H NMR spectrum of compound 6.

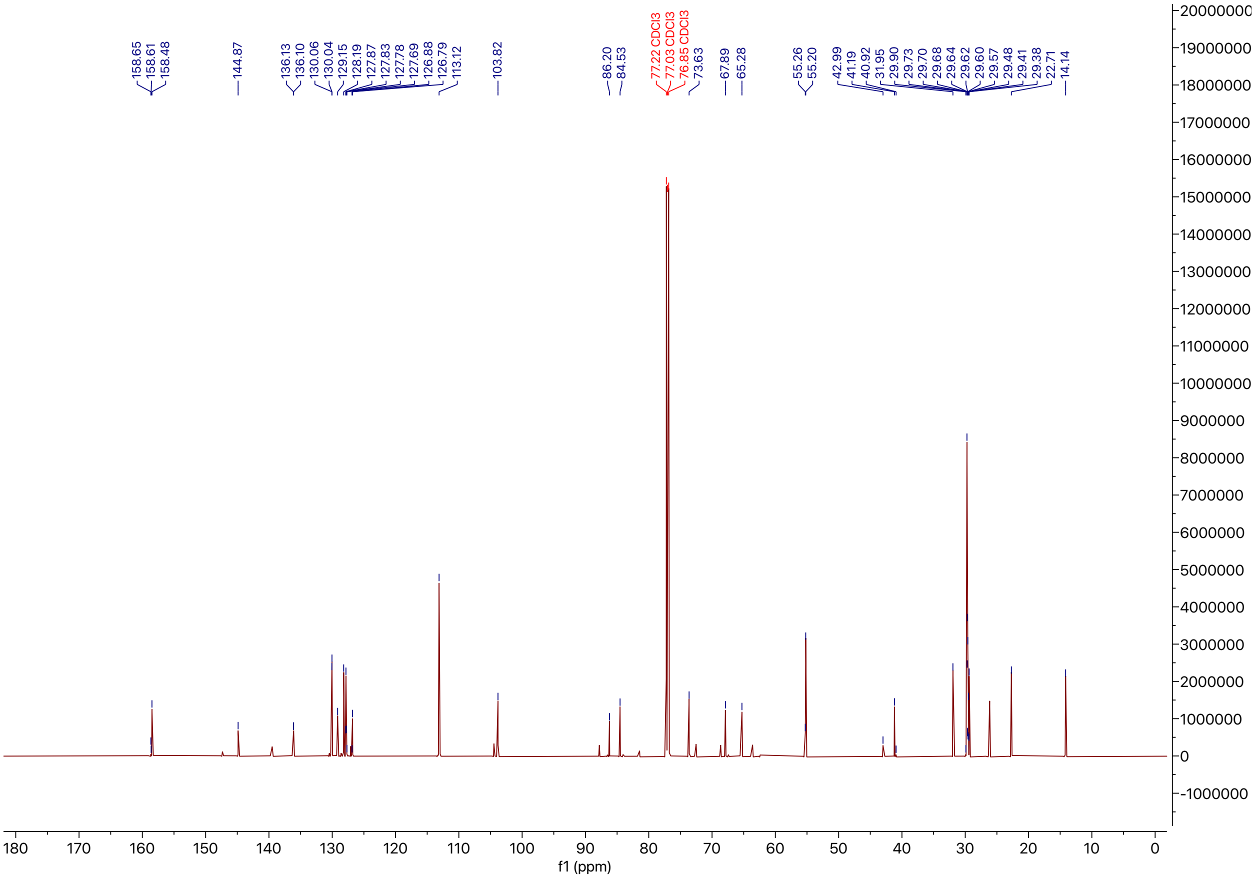

**Figure S23.** $^{13}$C NMR spectrum of compound 6.

**R-C18**

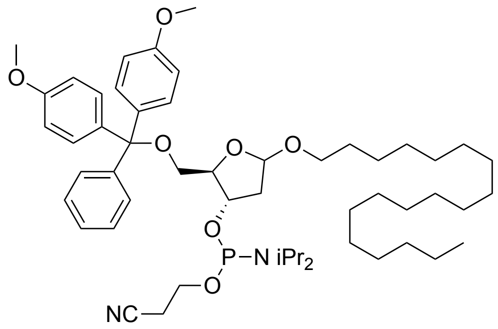

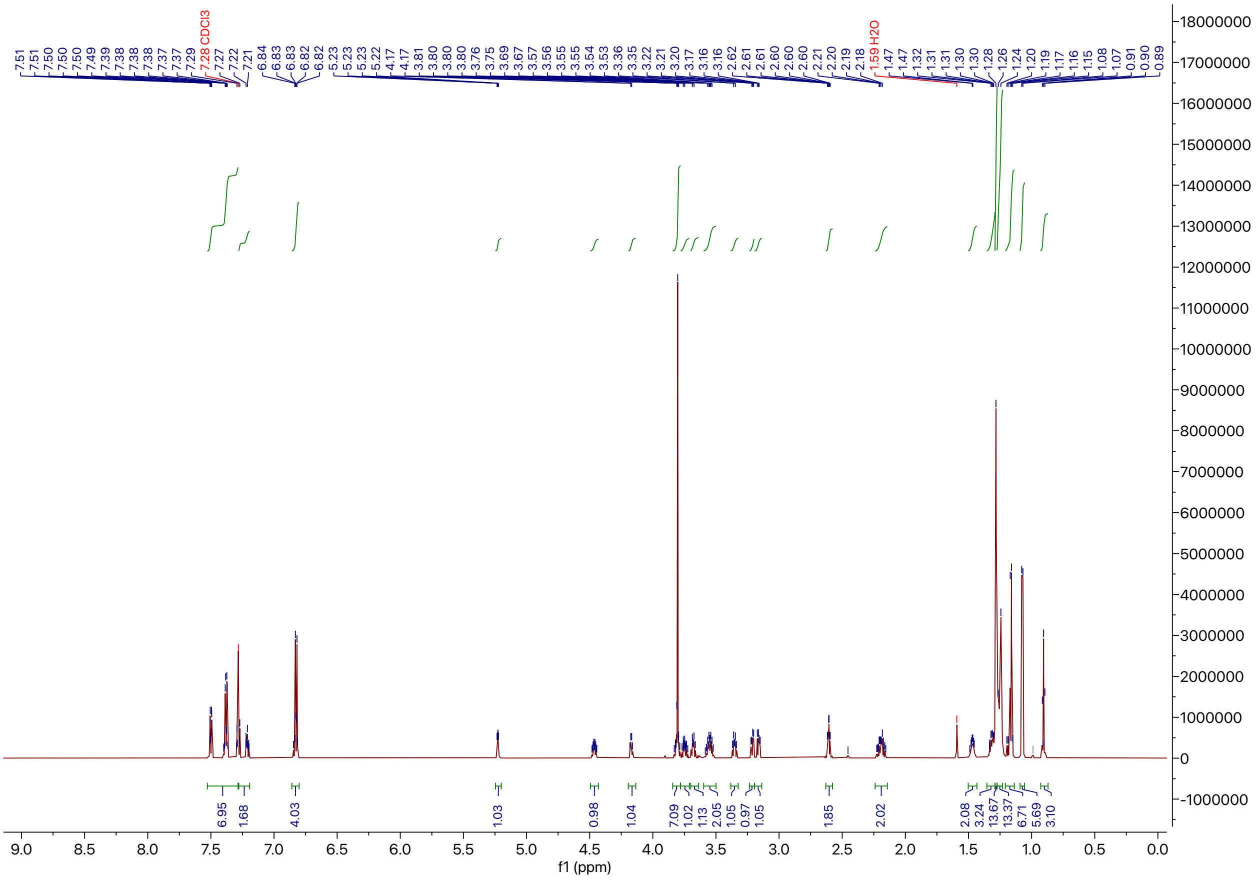

**Figure S24.** $^1$H NMR spectrum of R-$C_{18}$.

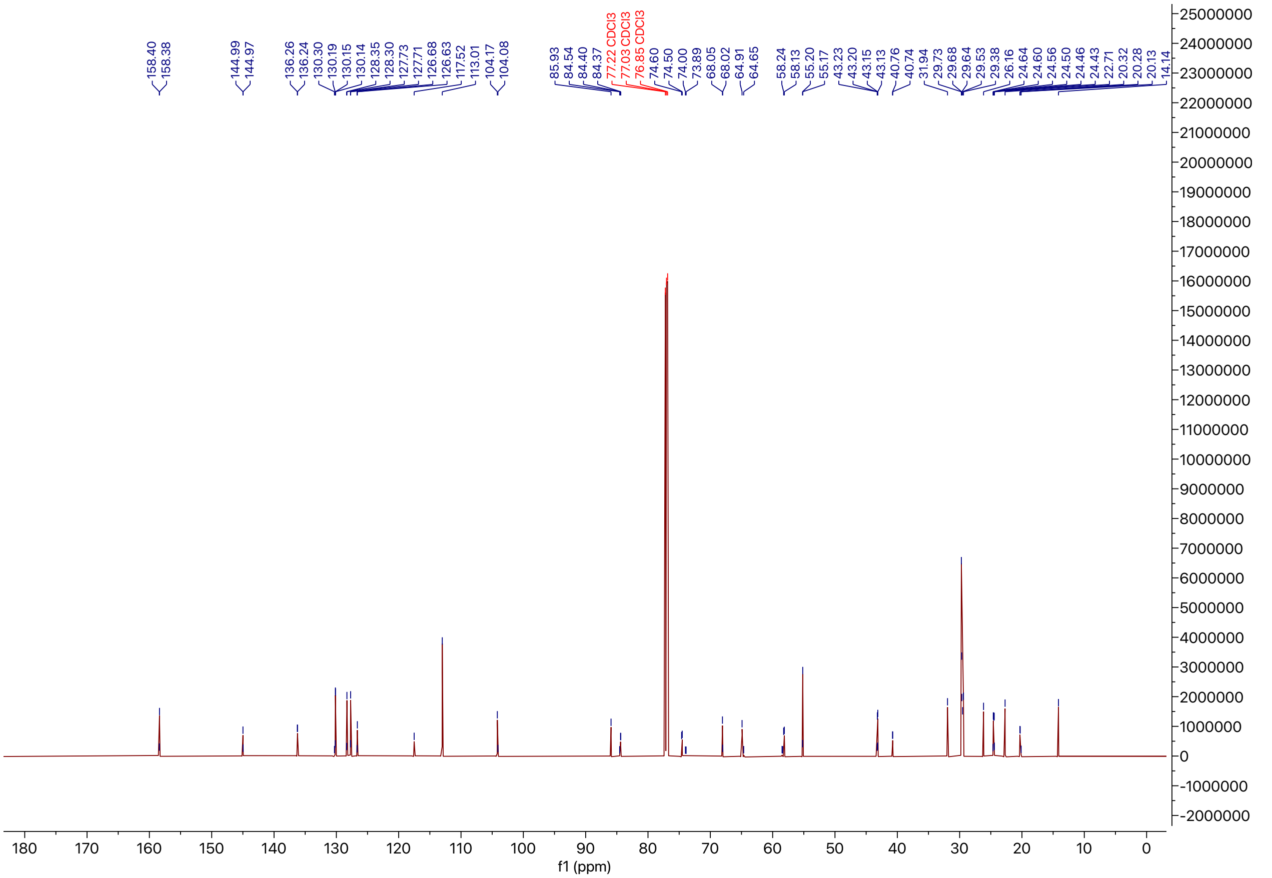

**Figure S25.** $^{13}$C NMR spectrum of R-$C_{18}$.
